# Glucagon-induced PGC-1α4/PPARγ promotes hepatic lipid storage and macrosteatosis in fasting and MASLD

**DOI:** 10.1101/2025.09.09.674670

**Authors:** Ayşim Güneş, Stewart Jeromson, Clémence Schmitt, Matthieu Schoumacher, James Eng, Aurèle Besse-Patin, Mélissa Léveillé, Laure Monteillet, Nathalie Jouvet, Cindy Baldwin, Isabelle R Frayne, Stephanie Petrillo, Anthoula Lazaris, Bich Nguyen, Matthieu Ruiz, Peter Metrakos, Jennifer L. Estall

## Abstract

The liver plays a central role in regulating the transition between fasting and feeding states, coordinating glycogen breakdown, gluconeogenesis, fatty acid catabolism, and lipid storage. Disruptions in this balance contribute to metabolic disorders, including Metabolic dysfunction Associated Steatotic Liver Disease (MASLD). Here, we identify a novel biological pathway downstream of glucagon that promotes the hepatic lipid accumulation associated with fasting. Using gain- and loss-of-function *in vivo* and *in vitro* models, we found that glucagon induces sustained expression of PGC-1α4, promoting lipid uptake and storage in hepatocytes by increasing PPARγ activity. Increased PPARγ/PGC-1α4 promotes hepatic *Fsp27/Cidec* expression, leading to lipid droplet expansion and triglyceride trapping in liver. Activity of the *PPARGC1A* alternative promoter and PGC-1α4 expression were higher in livers of patients with MASLD, and PGC-1α4 expression correlated with macrosteatosis. Consistently, persistent expression of hepatic PGC-1α4 in mice fed a western diet promoted macrosteatosis, exacerbated oxidative stress and altered hepatic lipid composition to resemble worsening Metabolic dysfunction Associated Steatohepatitis (MASH) in humans. Our findings demonstrate that glucagon-induced hepatic PGC-1α4/PPARγ activity facilitates efficient uptake and storage of lipids during fasting, but over-activation of this coordinated metabolic pathway leads to lipid accumulation and worsening of steatosis in MASLD.

## INTRODUCTION

The liver orchestrates an intricate transition between fasting and feeding states. Upon initiation of fasting, glucagon prompts the breakdown of glycogen and triglyceride stores. Concurrently, gluconeogenesis fosters synthesis of new glucose from non-carbohydrate precursors following glycogen depletion, and catabolism of fatty acids becomes a vital energy source. Prolonged fasting induces adipose tissue lipolysis, releasing free fatty acids that are subsequently taken up by the liver, where they are burned or stored to meet the continued energy demands of the body^1^. Hepatic fasting is not purely catabolic; instead, it represents a metabolic condition in which opposing anabolic and catabolic pathways operate in parallel. Catabolic oxidation provides fuel and energy for survival, while concurrent lipid uptake and storage ensures future nutrient availability and provides substrates for generation of alternative energy sources (e.g. ketones). Feeding, by contrast, constitutes a classic anabolic state, driven by insulin to suppress fasting-induced programs and promote nutrient storage. Proper coordination of these processes relies on transcriptional regulators functioning in harmony. Failure to maintain the delicate switch between these biological processes impairs energy balance and contributes to the pathogenesis of metabolic disorders, including (MASLD)^1, 2^.

In this study, we aimed to understand the roles and regulation of transcriptional proteins involved in the hepatic fasting response, in both physiological and pathological contexts. PPARγ and its coactivator PGC-1α are transcriptional regulators that orchestrate diverse metabolic processes across tissues, often exerting contrasting functions in the regulation of lipid metabolism and energy homeostasis. PPARγ is a transcription factor most known for its essential roles in adipose tissue, promoting the transcriptional program of adipogenesis. In liver, PPARγ has been shown to promote lipid uptake and *de novo* lipogenesis in the fed state, particularly following excessive caloric intake. It also inhibits inflammation, oxidative damage and endoplasmic reticulum stress^3^. Apart from the observation that its transcriptional expression declines in liver during fasting^4–6^, little is known about its regulation or function in this context.

In contrast, PGC-1α is induced in liver during fasting^7^, interacting with numerous transcription factors and transcriptional machinery to coordinate gluconeogenesis, beta oxidation and mitochondrial activity^8–10^. This coactivator has evolved as a critical node in the transition between fasting and fed states^11–14^. Beyond its metabolic role, PGC-1α exhibits both anti-oxidant^12^ and anti-inflammatory^15–17^ actions in liver to protect against metabolic stress. Interestingly, low levels of hepatic PGC-1α mRNA are reported in people living with obesity and MASLD^18–21^. Liver-specific ablation of the PGC-1α gene (*Ppargc1a*) in mice reduces hepatic lipid oxidation increasing steatosis, alters insulin signaling, increases oxidative stress and can drive fibrosis^11–13,21^. While PGC-1α was originally identified as a binding partner of PPARγ^22^, PPARγ is conversely increased in MASLD^23, 24^ and knock-out of hepatocyte PPARγ reduces steatosis in mice^25, 26^.

While their classical roles and differential expression patterns in physiological and pathological contexts imply a separation of PPARγ and PGC-1α function in liver, it is now appreciated that multiple PGC-1α proteins are transcribed from the *Ppargc1a* gene^27^. PGC-1α1 from the proximal promoter (canonical PGC-1α) drives the programs of mitochondrial oxidative metabolism and gluconeogenesis. Hepatocytes can also express NT-PGC-1α-a from the proximal promoter, PGC-1α-b and PGC-1α4 from the alternative promoter^16^, and L-PGC-1α from a liver-specific promoter^28^. NT-PGC-1α-a stimulates hepatic gluconeogenesis^29^ similar to the canonical protein, while PGC-1α4 increases expression of anti-apoptotic genes and suppresses cell death in response to TNFα and lipopolysaccharide^16^. Yet, strong induction of PGC-1α4 by glucagon in hepatocytes^16^ suggests an additional metabolic role that is not yet understood. In muscle, it has been shown that PGC-1α4 does not regulate mitochondrial or oxidative metabolism, but rather promotes myocyte hypertrophy and muscle strength^30^, increases glycolysis^31^, and prevents age-associated glucose intolerance and insulin resistance^32^, functions that are interestingly in line with the actions of PPARγ^33, 34^.

Using gain- and loss-of-function primary cell and mouse models to explore the rapid and dynamic shift between anabolic and catabolic states during fasting/re-feeding, we investigated how PGC-1α and its transcriptional partners coordinate the fasting response and impact development of steatotic liver disease in mice and humans living with MASLD. We found that hepatic PGC-1α4 is induced during fasting, where it interacts with PPARγ to stimulate *Fsp27/Cidec* expression, promoting lipid uptake and storage in hepatocytes. We propose that this function of PGC-1α4 ensures availability of fatty acids as a fuel substrate in liver during prolonged fasting. However, uncontrolled PGC-1α4 expression induces macrosteatosis in the liver, worsening liver pathology in the context of MASLD.

## MATERIALS AND METHODS

### Mice

Age- and sex-matched C57BL/6J male and female mice were maintained on ad-libitum chow (Teklad Global 18% Protein Rodent Diet) at 22°C (12h light-dark cycle). For gene and protein expression levels during fasting, 12-week-old wild-type (WT) male and female mice fed standard animal facility chow diet were fasted 24 hours. During the fasting to fed transition, 12-week-old WT^Cre+^ mice were fasted 16 hours and then re-fed 6 hours.

Hepatocyte-specific PGC-1α4 overexpressing male and female mice (PGC-1α4^HepTg+^)^16^ were compared to age- and sex-matched littermate Alb-Cre+ (WT^Cre+^) control mice, as well as PGC-1α4 transgene positive (WT^Tg+^) controls that have low, but detectible levels of ectopic PGC-1α4 transgene expression in all tissues^16^. Fasting was performed for 24 hours on mice aged 12 weeks. For re-feeding, mice were given access to food for 6 hours following a 16 hour fast. For MASLD modeling, mice were fed a high-fat, high-fructose, and cholesterol-rich diet (D17010103i, Research Diets, 20 kcal% protein, 40 kcal% carbohydrates, and 40 kcal% fat, with 2% w/w cholesterol) for 12 weeks, starting at 5 weeks of age.

PGC-1α4 shRNA (Integrated DNA Technologies) or control shRNA (shControl) was cloned into plasmid pAAV-shRNA-control purchased from Addgene (#85741). Adeno-associated virus stereotype 8 expressing the shRNA for PGC-1α4 (FWD 5’-AT AAA TGT GCC ATA TCT TCC A-3’) or the shControl (5’ -GGT-TCA-GAT-GTG-CGG-CGA-GT-3’) were purchased from Applied Biological Materials (ABM) Inc. Tail vein injections of either AAV-shControl or AAV-shPGC-1α4 viruses diluted in PBS (200 μl) were performed in 12-weeks old male mice at a dose of 1.5 x 10^11^ vector genomes per mouse (vg/mouse). Mice were monitored for 2-weeks post injection, after which the animals were fasted 24h prior to sacrifice and the livers collected. Successful knockdown of PGC-1α4 was confirmed by qPCR. All experiments were performed in accordance with IRCM animal facility institutional animal care and use committee regulations.

### Primary hepatocyte isolation

Primary hepatocytes were isolated from 12-weeks old male wild-type (WT) mice according to established protocols. Following over-night recovery in DMEM (Dulbecco’s Modified Eagle Medium) supplemented with 4.5 g/L glucose (Multicell, 319005-CL), 0.2% Bovine serum albumin (BSA) Fraction V, 2 mM sodium pyruvate, 1% penicillin-streptomycin, 100 nM dexamethasone, and 1 nM insulin, *Maintenance Media*), cells were subjected to a 16-hour incubation in low-glucose, insulin and dexamethasone-free media (DMEM, Multicell, 319-010-CL, supplemented with 1.0 g/L glucose, 0.2% BSA Fraction V, 2 mM sodium pyruvate, and 1% penicillin-streptomycin). For the glucagon time course experiment, cells were incubated with glucagon (50 nM, Cayman, 24204) for either 3, 6, 12, and 24 hours. For incubations longer than 12 hours, media was refreshed after 12 hours. For experiments with insulin, hepatocytes were treated with glucagon for 12 hours prior to addition of fresh media containing 50 nM glucagon and 100 nM insulin (Sigma, 16634) for an additional 12 hours.

For overexpression experiments, following overnight recovery in Maintenance Media, primary hepatocytes were infected overnight with adenoviral constructs expressing control (Ad-Vector), PGC-1α1 (Ad-PGC-1α1), or PGC-1α4 (Ad-PGC-1α4)^16, 30^ and incubated for a total of 48 hours in low-glucose, hormone-free media. To investigate roles for PPARγ, cells were pre-treated with a PPARγ antagonist (T0070907, Abcam, A13120876, 15 μM) for 2-3 hours, and/or treated with a PPARγ agonist (Rosiglitazone, sigma, R2408, 15 μM) in the presence and/or absence of a lipid mix (137.6 μM linoleic acid (Sigma, L1012), 75 μM oleic acid (Sigma, 0383), and 37.5 μM sodium palmitate (Sigma, P9767) for either 24 (RNA) or 48 hours (lipid content). For loss-of-function experiments, following overnight recovery in Maintenance Media, cells were exposed to adenovirus constructs expressing either a control sequence (shControl) or shRNA targeting PGC-1α4 mRNA (shPGC-1α4)^16, 30^ for 24 hours. Cells were then incubated with low-glucose hormone-free media for 16 hours, prior to treatment with lipids, agonists and/or antagonists, as above.

### RNA isolation, quantitative and non-quantitative PCR

RNA was isolated from approximately 100 mg of liver tissue using Trizol (Sigma, 15596018). cDNA synthesis (High-capacity reverse transcription kit, Applied Biosystems) was performed on 1 μg of RNA treated with DNAse-I and cDNA quantified using SYBR Green PCR mastermix (ABM, G891, G892). Data are presented as relative mRNA expression using the ΔΔCt threshold cycle method, with normalization to the Hypoxanthine-guanine phosphoribosyltransferase (Hprt) reference gene. To determine the presence or absence of full-length PGC-1α isoform mRNAs, non-quantitative PCR was performed using isoform-specific primers and a master mix (FroggaBio, FBTAQM). Detailed primer information is provided in **Table 1**.

### Protein isolation and western blotting

Total cell lysates were prepared using RIPA buffer and subjected to western blotting analysis using antibodies for anti-PGC-1α (Millipore, ST1202), anti-PPARγ (Santa cruz, 7196; 7273), anti-αHA (Cell signaling, 3724S), anti-FLAG (Millipore, F3165), HSP90 (Cell signaling, 4874S), anti-H3 (ABClonal, A2348), Anti-Tubulin (Cell signaling 2144S), and Beta actin (Sigma, A5441).

### Cellular fractionation and co-immunoprecipitation

For subcellular fractionation, cells were washed with cold PBS and resuspended in cytoplasmic lysis buffer (10 mM HEPES pH 7.5, 10 mM KCl, 3 mM MgCl2, 0.35 M sucrose, 0.1% NP40, 3 mM 2-mercaptoethanol, 0.4 mM phenylmethylsulfonyl fluoride, 1 μM pepstatin A, 1 μM leupeptin, and 5 μg/mL aprotinin). Following centrifugation, the supernatant (cytoplasmic protein) was saved isolated and the nuclear pellet resuspended in nuclear lysis buffer (3 mM EDTA, 0.2 mM EGTA, 1 mM dithiothreitol, 100 mM NaCl, and 0.8% NP40) and sonicated (30 s ON/30 s OFF) for 10 minutes. 1 mg of cytoplasmic or nuclear protein lysates were diluted in 1 mL of immunoprecipitation buffer (50 mM Tris, pH 8.0, 10 mM sodium pyrophosphate, 100 mM NaCl, 2 mM EDTA pH 8.0, 1% NP-40, and 10% glycerol, supplemented with protease/phosphatase inhibitors) and incubated with 2 μg of anti-HA antibody (Cell signaling, 3724S, to capture PPARγ) and protein A/G magnetic beads overnight at 4°C. The antibody-antigen complexes were eluted and subjected to immunoblot analysis using anti-FLAG antibody (Millipore, F3165, to detect FLAG-PGC-1α4).

### Oxygen consumption rate (OCR)

Primary hepatocytes (15,000 cells/well) were seeded onto 96-well plates. Following overnight recovery in Maintenance Media, hepatocytes were infected with indicated adenoviral constructs for 24 hours. Media was replaced with low-glucose hormone-free media for 6-8 hours (starvation) prior to and overnight incubation with the lipid mix (described above). 48 (over-expression experiments) or 72 (silencing experiments) hours post-transduction, cells were treated with 1 μM Oligomycin (Abcam, Ab141829,) or 1.6 μM FCCP (Abcam, Ab120081) in low-glucose hormone-free media. After 3 hours, FCCP wells were replenished with media supplemented with 1.0 μM rotenone (Abcam, Ab143145) for an additional 3 hours. DMSO served as the vehicle control. Plates were interfaced with the Resipher system (Lucid Scientific) for live, continuous monitoring of oxygen consumption. OCR values, calculated using Lucid Lab software, were normalized to total protein content of the cells and used to determine basal, coupled, spare, non-mitochondrial respiration, and proton leak values. Basal respiration was obtained by subtracting rotenone-treated OCR values from those of DMSO-treated control wells, while coupled respiration was determined as the difference between oligomycin- and DMSO-treated OCR values. Spare respiratory capacity was calculated by subtracting DMSO-treated values from those measured after FCCP treatment, and maximal respiratory capacity was defined as the difference between FCCP- and rotenone-treated OCR values. Proton leak was assessed as the difference between oligomycin- and rotenone-treated wells, whereas non-mitochondrial respiration was determined by subtracting OCR values of rotenone-treated wells from those of cell-free wells.

### Fatty acid oxidation

Primary hepatocytes were seeded into 12-well plates. Following overnight infection with adenoviral constructs in Maintenance Media, media replaced with low-glucose hormone-free media with or without 45 μM Etomoxir (Selleck chemicals, E1905) and incubated overnight. 48 (over-expression) or 72 (silencing) hours post-infection, cells were incubated with a master mix of non-bicarbonate assay buffer (114 mM of NaCl, 4.7 mM of KCl, 1.2 mM of KH_2_PO_4_, 1.2 mM of MgSO_4_, 0.5% fatty acid free BSA and 2.8-5 mM glucose) containing 0.9 μCi ^14^C-palmitate for 2 hours. Pre-wet (250 µl 2M NaOH) Whatman filter paper was used to capture labelled CO_2_ overnight following addition of 1.5ml 6N HCL to the media in sealed test tubes. Radioactivity of filter paper was measured using a beta counter for 1 minute.

### Glycerol release and total triglyceride measurement assays

Primary hepatocytes were seeded in 12-well plates. Following overnight infection with adenoviral constructs for 24 hours in Maintenance Media, cells were incubated with the lipid mix for 48 hours. Cells were then incubated with glucose-, pyruvate-, L-glutamine-, and phenol red-free media (Sigma D5030) supplemented with 0.2% fat-free BSA (GenDEPOT, A0100-010) for 2 hours to deplete glycogen. Media was then supplemented with 2-2.8 mM glucose, with or without isoproterenol (5-50 μM, Sigma I2760). After 3 hours, cell culture media was heated at 85°C for 10 minutes and centrifuged (5000 rpm) for 5 minutes. Free glycerol content in 120 μl media was measured (Free glycerol reagent, Sigma F6428). Total acylglycerol content was measured in cells following lysis in 2:1 chloroform-methanol and centrifugation (15000 rpm, 10 minutes at 4°C). The lower organic layer was collected, dried O\N and dissolved in a 60% Butanol:40% 2:1 Triton X-114:Methanol buffer, and total acylglycerols measured using a kit (Thermo Scientific, TR22421).

### Bodipy staining

Cells were seeded onto 15 mm cover slips pre-coated with poly-L-lysine (Sigma, P6282); and collagen I (0.3 mg/ml) (Corning, 354236). Cells were incubated with Bodipy (Sigma, D3835) diluted in culture medium at 5 μM for 30 minutes. Medium was aspirated and cells fixed with 4% paraformaldehyde in PBS for 15 minutes at 37°C. Cells were washed three times with PBS and cover slips mounted using DAPI containing mounting media (Invitrogen, P36935). Imaging was performed using (DM4000 MLED Versatile upright microscope, Leica). Bodipy fluorescence was excited at 558/568 nm with emission fluorescence detected using an internal Hybrid (HyD) detector. Background signals in representative images were reduced by calculating/subtracting strategies (Image J).

### Hepatic lipid, glycogen, glucose and G6P

Lipid tissues were homogenized with zirconium beads in cold PBS for 3 min. Homogenates were resuspended in a 2:1 chloroform and methanol solution and lipids isolated and measured as described above. Total acylglycerol content was quantified by Total Triglyceride Reagent Kit (Thermo Scientific, TR22421).

Liver was homogenized in 8 volumes (w/v) of 6% PCA with zircon 0.5 mm beads for 3 min. Following centrifugation at 10 000 rpm at 4C for 15 min, the supernatant was neutralized with 1/10 volume of K_2_CO_3_ and pH was adjusted in between 6.5 and 8.5. After 1 hour on ice, tubes were centrifuged at 10 000 rpm (4°C, 5 min) and supernatant, by using the Keppler and Decker method as previously described^35^. The supernatant collected was used to measure G6P, glucose and glycogen. To measure G6P, which represents endogeneous phosphorylated glucose pool, ½ volume of NaOH (0.3M) was added and samples boiled for 20 min. 6 μl of sample was mixed with 100 μl of assay-mix 1 (TEA Buffer 0.3M pH7.5, ATP/NADP and G6PDH 50U/ml) in one well of 96-well plate and measured at 340 nm. To measure free glycose, these wells were incubated with HK (37.5 U/ml per well) to phosphorylate glucose at room temperature for 30 min and then measured at 340nm. To measure glycogen, stored glycogen was denatured by incubation with 1.5 volumes of freshly prepared α-Amyloglucosidase (1mg/ml) for 1h at 45°C. For G6P, 20 μl of supernatant was mixed with 80 μl of assay-mix 2 (TEA Buffer 0.3 M pH7.5 and ATP/NADP) to the well and measured at 340 nm, Following the additions of HK (37.5 U/ml) and G6PDH (50 U/ml per well), and the tubes were incubated room temperature for 30 min to measure the OD at 340 nm. Glycogen content was calculated from the difference between amyloglucosidase-treated and untreated samples. Absorbance was measured at 340 nm before and after enzyme addition, and metabolite concentrations were calculated from standard curves prepared with glucose and G6P. Results were normalized to tissue weight (mg glycogen per g tissue, or µmol glucose or G6P per g tissue).

### Serum lipids and alanine aminotransferase

Total acylglycerols were measured in 2 μl serum using a commercial kit (Thermo Scientific, TR22421). Alanine aminotransferase (ALT) was measured using the Liquid ALT (SGPT) Reagent Set (Pointe Scientific Inc., A7526) as per manufacture’s instructions.

### Magnetic resonance imaging

Live animals were weighed and ear-tags removed before magnetic resonance imaging (MRI) using the minispec mq 7.5 NMR analyzer (Bruker, MA). Lean and fat mass was measured according to manufacturer’s instructions.

### Pyruvate tolerance test

After an overnight fast, mice were weighed and injected intraperitoneally (i.p.) with freshly prepared sodium pyruvate dissolved in sterile PBS (pH adjusted to 7.4) at a dose of (2 g/kg body weight). Blood glucose levels were measured from the tail vein at baseline (0 min) and at 15, 30, 45, 60, 75 and 90 minutes after injection using a handheld glucometer.

### Histology and scoring

Liver tissue was fixed in 10% formalin, paraffin-embedded, sectioned at 3-5 μm, and stained with hematoxylin and eosin and Masson’s trichrome stain. A pathologist, blinded to sample identity, evaluated simple steatosis, lobular inflammation, and fibrosis according to the FLIP algorithm as described by Bedossa et al.^36^ Detailed scoring methodology has been reported previously^12^.

### Untargeted LC-MS Lipidomic Analysis

Liver samples were harvested from male and female PGC-1α4^HepTg+^ and WT^Cre+^ mice (n = 10-12 per group) fed a MASLD-promoting diet, as described above. Untargeted lipidomic analyses were performed using validated protocols for lipid extraction and LC-MS analysis, as previously described^37, 38^ with slight modifications.

Briefly, lipids were extracted from 30 mg of liver tissue and spiked with six internal standards (PS(12:0/12:0), PC(14:0/14:0), PE(17:0/17:0), PC(19:0/19:0), Cer(d18:1/10:0), and TG(17:0/17:1/17:0)-d5). Samples were injected (volume=0.5 µL) into a 1290 Infinity HPLC coupled to a 6550 QTOF equipped with a dual ESI source (Agilent Technologies Inc., Santa Clara, USA) and analyzed in positive mode. Lipids were eluted on a Zorbax Eclipse plus C18, 2.1 x 100 mm, 1.8 µm (Agilent Technologies Inc.) heated to 40°C with a constant flow rate at 0.45 mL/min with an 83 min gradient of mobile phase A (0.2% formic acid and 10 mM ammonium formate in water) and mobile phase B (0.2% formic acid and 5mM ammonium formate in methanol/acetonitrile/MTBE 55:35:10 (v/v/v)). Data collection and processing were done using a previously validated in-house pipeline^38^ including (1) application of a frequency filter of 80% based on a feature’s presence within each condition, (2) data normalization using cyclic loess algorithm (from the *limma*^39^ R package from Bioconductor), (3) imputation of missing values to 90% of the lowest value, and (4) batch correction for manipulator using *ComBat*^40^. The final dataset consisted of a matrix in which each sample is described by 1778 features.

For statistical analysis, Principal Component Analysis (PCA) was performed using *mixOmics* R package^41^ for data overview and outlier identification. Group comparisons were made through linear regression analysis, p-values were adjusted using False Discovery Rate (FDR) and corrected for sex (*limma*^39^ R package). Most discriminant features between the two groups were chosen using an adjusted p-value <1×10e-6 and a fold-change (FC) >1.25 or <0.8 threshold. Annotation was done using in-house database (based on exact mass, retention time, and MS/MS scans) and public LIPID MAPS database with further validation using MS/MS analysis for the most discriminant features.

### Human liver samples

Liver tissue was collected from patients undergoing hepatic resection at the McGill University Health Centre after informed consent was obtained and stored as part of the McGill Liver Disease Biobank. Subjects were included if they had a diagnosis of MASLD or MASH (absence of known viral infection), or low or absent steatosis (control livers), based on pathologist scoring of liver histology using NAFLD Activity Score (NAS): Low =<2, NAFLD =3-5, NASH =6-9 and fibrosis staging from 1A to 3. Forty-nine subjects aged 33-81 years fit our criteria (Healthy / Low steatosis n=15, MAFL n=20, MASH n=14). Samples were snap-frozen and stored at −80°C. Study protocol was approved by the Research Ethics Boards of McGill and the Institut de recherches cliniques de Montréal (IRCM).

### Statistical analysis

Outliers were identified, and normality tests (Shapiro–Wilk test) performed. An unpaired *t*-test (normally distributed data) or a non-parametric Mann–Whitney *U* test was applied when comparing two variables. For multiple measurements of the same sample, multiple unpaired *t*-tests or Mann–Whitney *U* tests were conducted with correction for multiple comparisons. When comparing three experimental groups with one variable, either a one-way ANOVA or a Kruskal-Wallis test was employed. For experiments with two variables, a two-way ANOVA was performed followed by Post hoc analyses, as indicated. For two-way ANOVA, statistical significance was assessed by a Sidak’s multiple comparisons post hoc test. For *in vitro* analyses, representative data from 3–4 independent experiments are shown. Data are presented as mean ± standard deviation (SD) for *in vitro* experiments,and mean ± standard error of the mean (SEM) for *in vivo* experiments. All statistical analyses were conducted using GraphPad Prism software.

## RESULTS

### PGC-1α4 protein levels persist in the liver during fasting and are repressed by refeeding

We hypothesized that activity of multiple PGC-1α proteins during fasting coordinates the balance between fuel partitioning, catabolism and nutrient storage based on metabolic demand. Canonical *Pgc-1α1* is induced in liver by fasting and repressed by re-feeding^42, 43^. We found this was also true for hepatic *Pgc-1α4* mRNA in male (**Figure 1A,B**) and female **(Supplementary Figure 1A,B**) mice. Consistently, both *Pgc-1α1* and *Pgc-1α4* mRNA were induced in primary hepatocytes treated with glucagon, and this was suppressed by insulin (**Figure 1C**). *In vitro*, glucagon-induced *Pgc-1α1* and *Pgc-1α4* mRNA peaked at 3 hours and again after 24 hours (**Figure 1D**); however, only protein levels for PGC-1α1 mirrored this temporal pattern (**Figure 1E,F**). In contrast, PGC-1α4 protein peaked at 6 hours and was maintained over the next 18 hours (**Figure 1E,F**), consistent with enhanced PGC-1α4 protein stability^44^. These results indicate that while various *Ppargc1a* promoters respond similarly to fasting and refeeding hormonal cues, PGC-1α isoforms proteins have divergent expression patterns.

**FIGURE 1.**
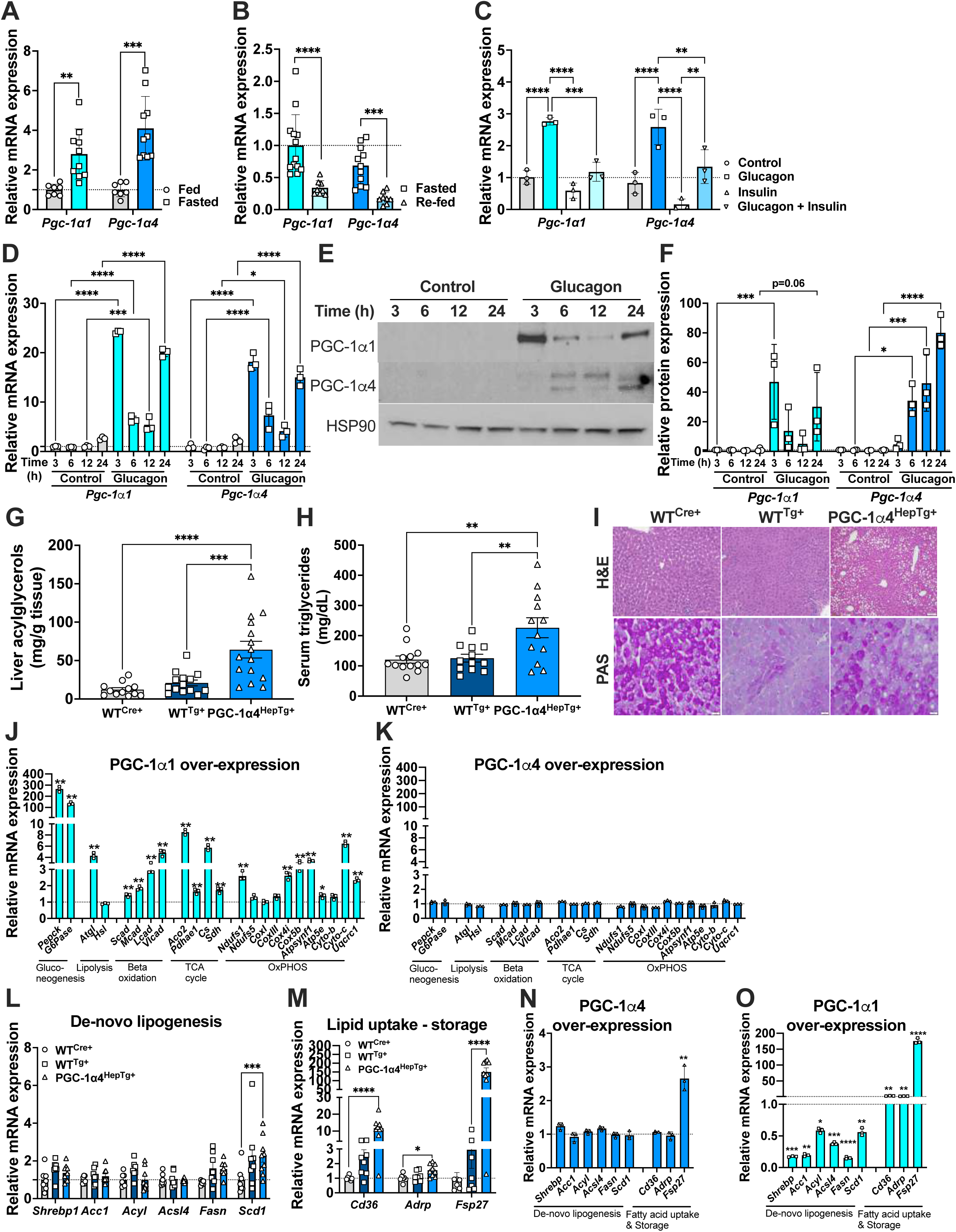
PGC-1α1 and PGC-1α4 are induced by fasting and repressed by feeding in mouse liver. Relative hepatic mRNA levels in the livers of 12-week-old C57BL/6J male mice **A)** fasted for 24 hrs (n=7-10) or **B)** fasted for 16 hrs and re-fed for 6 hrs (n=8-13). **(C)** Relative mRNA levels in primary hepatocytes treated with 50 nM glucagon, 100 nM insulin, or the combination for 12 hours (n=3). **D)** mRNA and **E,F)** protein expression of PGC-1α1 (∼115kDa) and PGC-1α4 (∼38kDa) in primary hepatocytes treated with 50 nM glucagon (n=2). **(G)** Total liver acylglycerols, **(H)** serum total triglycerides, and **(I)** representative H&E and PAS staining of liver sections from 8-12 weeks old male mice (n=7-15). **(J,K,N,O)** Relative mRNA expression in primary hepatocytes (n=2-4 independent experiments done in triplicate). **(L,M)** Relative mRNA expression in the livers of 12-week-old male mice (n=7-9). Bars represent means ± SEM for *in vivo* and ± SD for *in vitro* data, *P ≤ 0.05, **P ≤ 0.01, ***P ≤ 0.001, ****P ≤ 0.0001.

Surprisingly, livers of young, *ad libitum*-fed male and female mice overexpressing PGC-1α4 in hepatocytes (PGC-1α4^HepTg+^) had significant hepatic steatosis associated with increased liver and serum acylglycerols (**Figure 1G-I, Supplementary Figure 1C-E**) when compared to littermate control mice carrying either Albumin-Cre alone (WT^Cre+^) or the (Lox-stop-Lox) PGC-1α4 transgene alone (WT^Tg+^). This is in sharp contrast to mice with liver-specific canonical PGC-1α1 over-expression, which have low hepatic lipids due to increased fat oxidation^10^. Of note, while hepatic and serum lipids were high, hepatic glycogen was low in PGC-1α4^HepTg+^ livers (**Figure 1I, Supplementary Figure 1E**). Thus, PGC-1α4 appears to have a metabolic role quite distinct from PGC-1α1.

### PGC-1α4 does not regulate the same gene program as PGC-1α1

Canonical PGC-1α1 drives expression of genes associated with glucose metabolism, lipid catabolism, mitochondrial activity, and detoxification of reactive oxygen species^9, 42, 45, 46^. To shed light on the metabolic processes controlled by PGC-1α4, gene levels were measured in primary hepatocytes following PGC-1α1 or PGC-1α4 overexpression **(Supplementary Figure 2A**). As expected, PGC-1α1 significantly increased mRNAs associated with gluconeogenesis, lipolysis, beta-oxidation, TCA cycle and oxidative phosphorylation (**Figure 1J**). Notably, none of these genes were influenced by PGC-1α4 overexpression (**Figure 1K).** Thus, despite having >95% amino acid identity with the N-terminus of canonical PGC-1α1 and expressing the full ‘activation’ domain, PGC-1α1 and PGC-1α4 share very few metabolic gene targets in hepatocytes.

Since hepatic lipid levels were high in PGC-1α4^HepTg+^ mice, we explored whether pathways of *de novo* lipogenesis, lipid uptake and/or lipid storage were altered *in vivo*. Livers of PGC-1α4^HepTg+^ mice had significantly increased *Scd1* (fatty acid synthesis) (**Figure 1L, Supplementary Figure 1F**), along with *Cd36, Adrp* and *Fsp27/Cidec* (lipid uptake and storage) (**Figure 1M, Supplementary Figure 1G**). However, acute PGC-1α4 overexpression in primary hepatocytes did not impact mRNAs of genes controlling *de novo* lipogenesis or lipid uptake (**Figure 1N**), in contrast to PGC-1α1, which suppressed and induced these pathways, respectively (**Figure 1O**). Interestingly, both PGC-1α1 and PGC-1α4 significantly increased expression of *Fsp27/Cidec* (an adipocyte lipid droplet fusion/expansion protein) (**Figure 1N,O**).

Notably, *Scd1*, *Cd36*, *Adrp* and *Fsp27/Cidec* are well-established transcriptional targets of PPARγ, a nuclear receptor that plays a central role in promoting lipid storage and adipogenic gene expression. While overexpression of PGC-1α4 was not sufficient to induce most PPARγ targets (**Figure 1N**), silencing of this specific PGC-1α isoform **(Supplementary Figure 2B, 3A)** significantly blunted *PPARγ*, *Cd36*, *Fsp27* and *Scd1* mRNA **(Supplementary Figure 3B-F**). Taken together with *in vivo* data, disruption of hepatic PGC-1α4 seems to compromise hepatic PPARγ target genes, but additional cellular signals are likely needed to facilitate coactivation. Given the consistent and robust regulation of *Fsp27* by PGC-1α4, and the corresponding macrosteatosis in PGC-1α4^HepTg+^ livers (**Figure 1I, Supplementary Figure 1E**), we focused on the biological significance of the relationship between PGC-1α4, PPARγ and FSP27 during fasting.

### PGC-1α4 interacts with PPARγ to drive *Fsp27* expression during fasting

PPARγ activity in hepatic fasting may seem confounding, as most published studies show that PPARγ expression decreases during fasting^4–6^. However, nearly all of studies report only mRNA levels. Consistently, we also observed that *Pparγ* mRNA levels were reduced in livers of fasting male mice **(Supplementary Figure 4A,B**), as well as in primary hepatocytes treated with glucagon **(Supplementary Figure 4C**). However, reduced *Pparγ* mRNA was also observed upon refeeding male mice **(Supplementary Figure 4D,E**), and following insulin treatment in primary hepatocytes **(Supplementary Figure 4C**). Importantly, hepatic PPARγ protein was induced in fasted male and female mice, and decreased following re-feeding (**Figure 2A,B**), parallelling expression patterns of PGC-1α4 (**Figure 1A,B; Supplementary Figure 1A,B**), and *Fsp27* mRNA (**Figure 2C,D**). In primary hepatocytes, glucagon significantly induced PPARγ protein and *Fsp27* mRNA, while insulin repressed glucagon induced expression of PPARγ and *Fsp27* (**Figure 2E,F**), indicating increased PPARγ activity in hepatic fasting.

**FIGURE 2.**
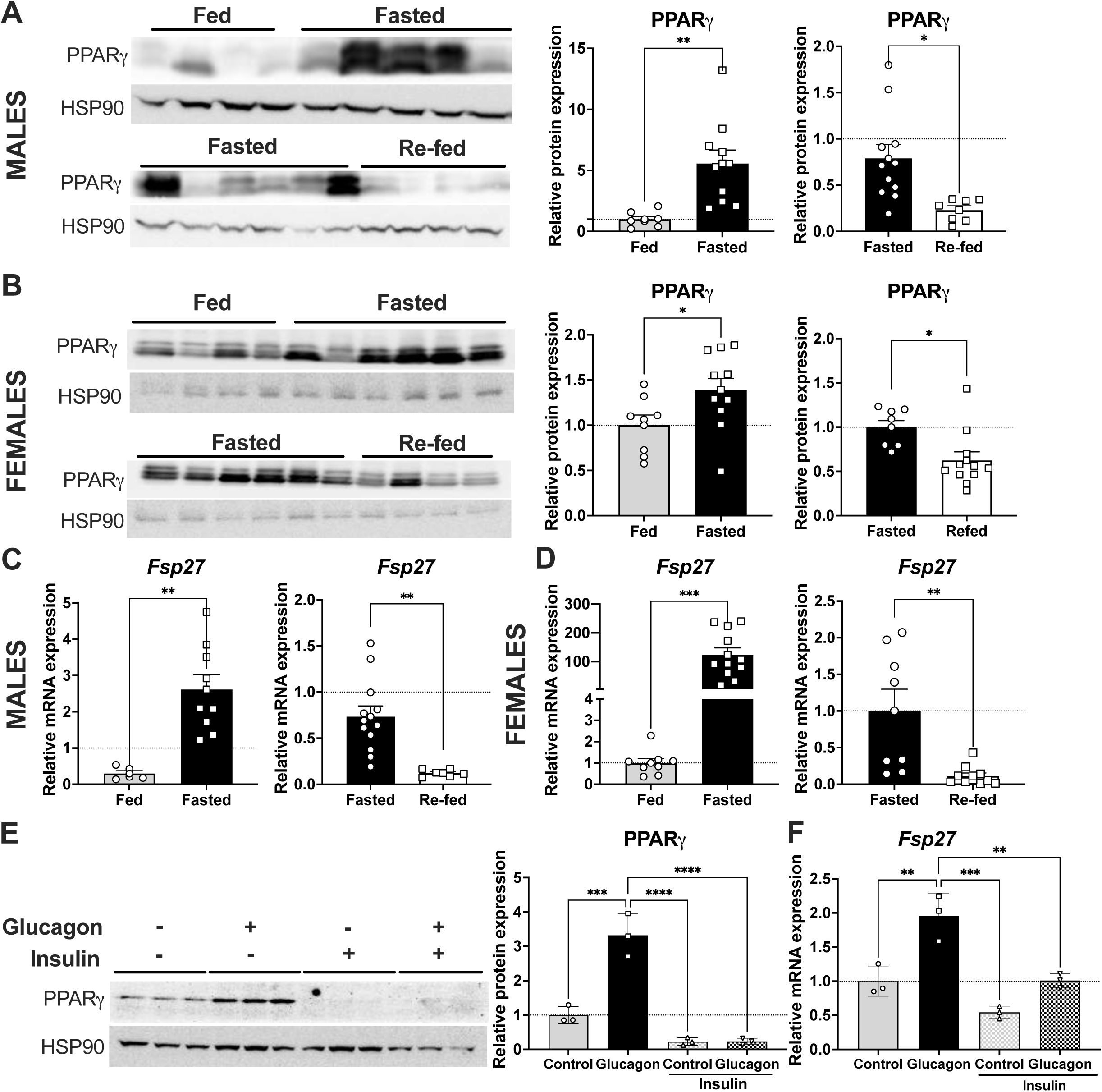
Fasting induces and re-feeding represses hepatic PPARγ and Fsp27 expression. Relative protein expression of PPARγ in livers of 12-week-old C57BL/6J **(A)** male and **(B)** female mice after 24 hrs of fasting. Relative *Fsp27* mRNA levels in livers of **(C)** male and **(D)** female C57BL/6J (WT^Cre+^) mice re-fed for 6 hrs following a 16-hr fast. Data are expressed as means ± SEM (n=7-12). **(E)** Relative protein and **(F)** mRNA levels in primary hepatocytes treated with 50 nM glucagon for 24 hrs +/- 100 nM insulin for the last 12 hrs. Data are mean ± SD representative of 2-3 independent experiments done in triplicate. *P ≤ 0.05, **P ≤ 0.01, ***P ≤ 0.001, and ****P ≤ 0.0001.

As PGC-1α4 can coactivate PPARγ^47^, we hypothesized that PGC-1α4 might interact with PPARγ in liver, and this interaction might play a role in the regulation of *Fsp27* during fasting. We found that PGC-1α4 physically interacted with PPARγ in both the cytoplasm and nucleus of hepatocytes, and the nuclear PGC-1α4-PPARγ interaction increased following exposure of hepatocytes to a physiological mix of free fatty acids (**Figure 3A**). This suggests that lipid signals (which can act as receptor ligands for PPARγ) may drive translocation or stabilization of the PGC-1α4-PPARγ complex in the nucleus, facilitating target gene transcription.

**FIGURE 3.**
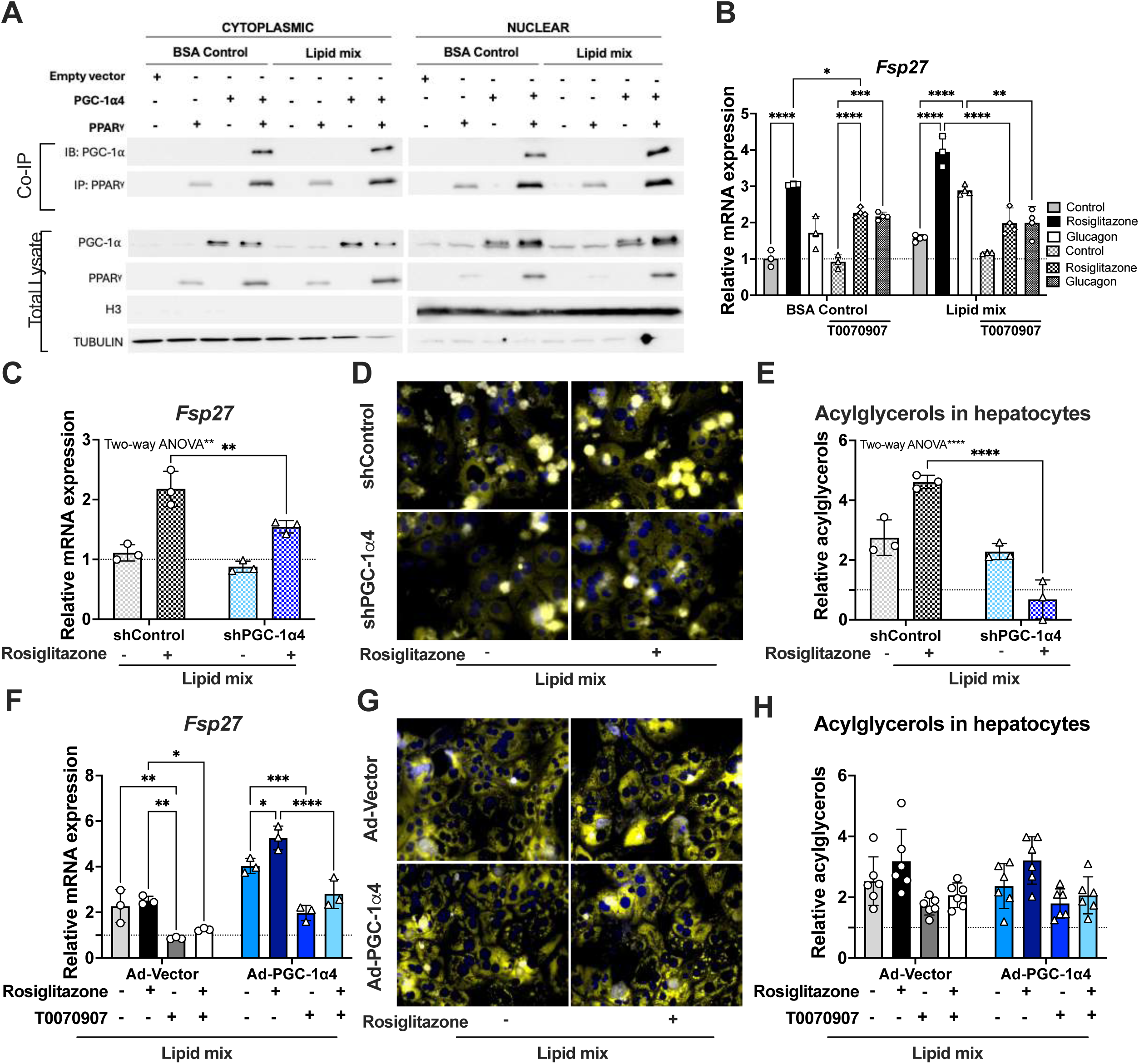
PGC-1α4 interacts with PPARγ to induce Fsp27 and lipid storage in hepatocytes. **(A)** Protein expression measured by Western blot following co-immunoprecipitation from cytoplasmic or nuclear protein lysates from primary hepatocytes over-expressing PGC-1α4 and PPARγ treated with or without a mix of free fatty acids (lipid mix: 137.6 µM linoleic acid, 75 µM oleic acid, and 37.5 µM sodium palmitate) for 36 hrs. **(B)** Relative mRNA expression in primary hepatocytes pre-treated with or without a PPARγ antagonist (T0070907, 15 µM), prior to incubation with 50 nM glucagon or 15 µM rosiglitazone, in the presence or absence of the lipid mix. **(C)** Relative mRNA expression, **(D)** Bodipy staining of lipid droplets and **(E)** acylglycerols levels in primary hepatocytes expressing shPGC-1α4 or shControl treated with rosiglitazone and the lipid mix.c**(F)** Relative mRNA expression, **(G)** Bodipy staining of lipid droplets and **(H)** acylglycerols levels in primary hepatocytes expressing Ad-PGC-1α4 or Ad-Vector control pre-treated with T0070907 prior to incubation (as indicated) with rosiglitazone and the lipid mix. Bars are mean ± SD of triplicate replicates of a representative experiment from 2-4 independent experiments. \**P* ≤ 0.05, \*\**P* ≤ 0.01, \*\*\**P* ≤ 0.001, \*\*\*\**P* ≤ 0.0001.

Hepatic *Fsp27* is increased during fasting^4, 48^ and can promote lipid storage^5^. However, it remains an open question which biological stimuli (e.g. hormones, metabolites) and transcription factors control *Fsp27* expression. Previous studies have shown that *Fsp27* transcription can be controlled by PPARγ and/or cyclic AMP-responsive element-binding protein (CREB)^5, 48–50^. Considering the strong physical interaction between PGC-1α4 and PPARγ, we investigated PPARγ’s role on *Fsp27* expression within the context of fasting, both in the presence and absence of lipid signals. Lipids alone raised *Fsp27* levels and enhanced increases associated with glucagon or PPARγ agonist (Rosiglitazone) ((**Figure 3B**). Furthermore, the PPARγ antagonist (T0070907) only blunted glucagon-induced *Fsp27* when lipids were present, demonstrating that the increase in *Fsp27* downstream of glucagon was dependent on both PPARγ and lipid signals.

Next, we investigated whether PGC-1α4 is important for PPARγ-mediated *Fsp27* transcription and lipid storage. Rosiglitazone and glucagon-induced *Fsp27* was significantly attenuated by knockdown of endogenous PGC-1α4 in primary hepatocytes (**Figure 3C, Supplementary Figure 5A-C**). In parallel, accumulation of intracellular lipids following rosiglitazone treatment was completely abolished by PGC-1α4 silencing (**Figure 3D,E**). PGC-1α4-induced *Fsp27* transcription was also blocked by antagonism of PPARγ (**Figure 3F, Supplementary Figure 5D-F**). However, overexpression of PGC-1α4 alone did not promote accumulation of intracellular acylglycerols (**Figure 3G,H**). Collectively, our data indicate that PGC-1α4 enhances PPARγ-dependent *Fsp27* transcription downstream of glucagon, and is necessary but not sufficient for hepatic lipid accumulation in hepatocytes.

### PGC-1α4 prevents the hydrolysis of hepatic triglycerides and decreases fatty acid oxidation

In addition to expanding lipid droplet size, hepatic FSP27 overexpression reduces fatty acid release and inhibits beta oxidation^51^. Thus, we hypothesized that activation of the PGC-1α4/PPARγ/FSP27 axis during fasting serves to sequester lipids, limiting accessibility for oxidation. During fasting, adrenergic signaling mobilizes hepatic energy stores^52^. Consistently, isoproterenol stimulated hepatic lipolysis, causing a dose-dependent increase in free glycerol (**Figure 4A**) while decreasing stored acylglycerol (**Figure 4B**). PGC-1α4 overexpression led to markedly lower basal and isoproterenol-stimulated glycerol release (**Figure 4A**) and prevented isoproterenol-induced triglyceride breakdown (**Figure 4B**). Conversely, silencing of PGC-1α4 increased lipolytic glycerol release (**Figure 4C**) and significantly reduced intracellular triglycerides stored during lipid loading (**Figure 4D**), demonstrating the importance of PGC-1α4 in maintaining hepatic acylglycerol levels. In contrast, canonical PGC-1α1 increased glycerol release (**Figure 4A**) and lowered hepatic triglycerides (**Figure 4B**), consistent with its known role in lipid catabolism^10, 13^. Mirroring FSP27, PGC-1α4 reduced oxidation of ^14^C-palmitate compared to control cells (**Figure 4E**), in stark contrast to the fatty acid oxidation-promoting effects of PGC-1α1. Silencing PGC-1α4 increased fatty acid oxidation (**Figure 4F**). These data highlight the lipid storage properties of PGC-1α4 versus the catabolic activity of PGC-1α1. Lastly, the observed effects on lipid metabolism were not explained by differential regulation of oxidative phosphorylation, as PGC-1α1 and PGC-1α4 had similar effects on cellular respiration (**Supplementary Figure 6**). In light of increased lipid storage, decreased lipolysis and prevention of fatty acid oxidation, we propose that the increased hepatic PGC-1α4/PPARγ/FSP27 axis during fasting promotes effective storage of lipids.

**FIGURE 4.**
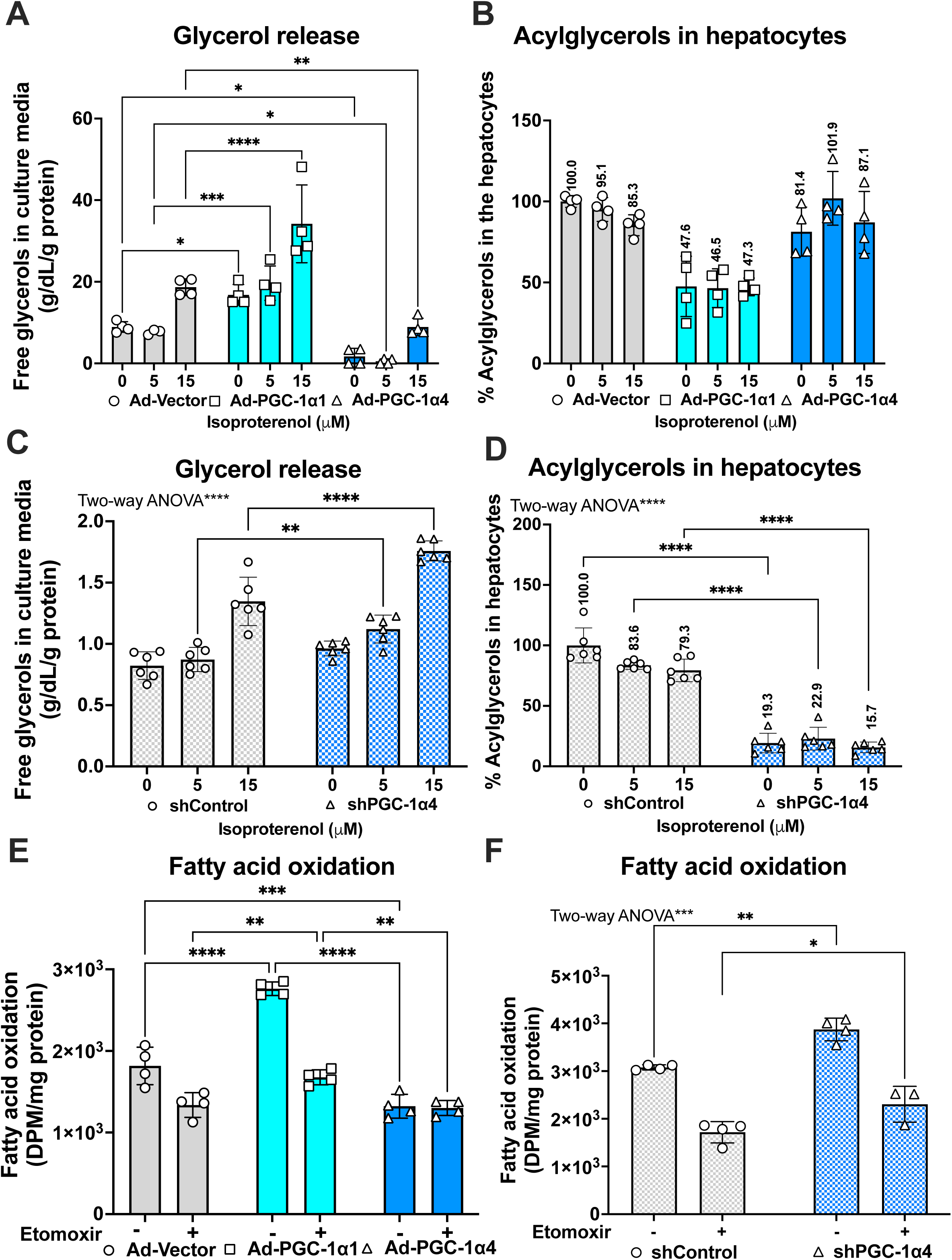
PGC-1α4 attenuates isoproterenol-induced lipolysis and lipid oxidation in primary hepatocytes. **(A,C)** Free glycerol concentration in cell culture media and **(B,D)** acylglycerol levels in primary hepatocytes expressing either Ad-PGC-1α1, or Ad-PGC-1α4, shPGC-1α1, shPGC-1α4) or controls (Ad-Vector, shControl). Cells were pretreated with the lipid mix prior to glycogen depletion and then treatment with isoproterenol for 3 hrs (n=4-5). **(E,F)** Fatty acid oxidation of C^14^-palmitate (n=4). Bars represent the mean ± SD of triplicate replicates from one experiment representative of 4-5 independent tests. *P ≤ 0.05, **P ≤ 0.01, ***P ≤ 0.001, and ****P ≤ 0.0001.

### Prolonged expression of PGC-1α4 causes macrosteatosis in liver

From a physiological perspective, we proposed that this mechanism helps provide a lasting source of lipid energy for use during prolonged nutrient deprivation (starvation). To test this, we fasted PGC-1α4^HepTg+^ mice and controls for 24 hours. Fasted PGC-1α4^HepTg+^ mice maintained higher hepatic acylglycerols (**Figure 5A, Supplementary Figure 7A**), had larger livers (**Figure 5B,C, Supplementary Figure 7B,C**) and high serum triglycerides (**Figure 5D, Supplementary Figure 7D**). Notably, livers of male and female PGC-1α4^HepTg+^ mice had predominant macrosteatosis (**Figure 5E, Supplementary Figure 7E**), correlating with higher PPARγ and *Fsp27* expression (**Figure 5F-H, Supplementary Figure 7F-H**).

**FIGURE 5.**
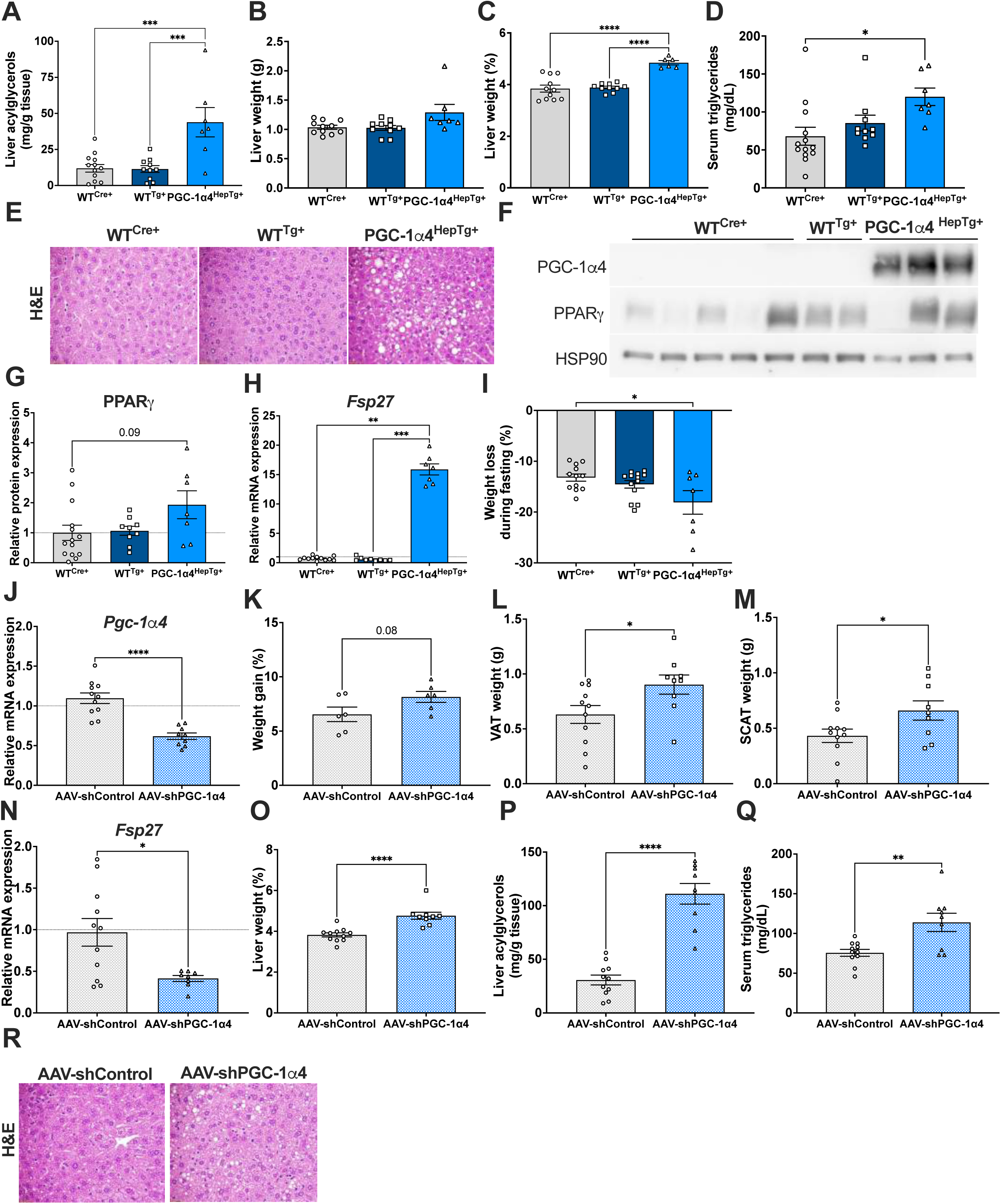
Prolonged expression of PGC-1α4 stimulates macrosteatosis in liver. **(A)** Liver acylglycerol levels, **(B,C)** liver weights, **(D)** serum triglyceride levels, **(E)** representative images of H&E stained of liver sections, **(F)** liver protein expression measured by Western blot **(G)** quantified by densitometry, and **H)** relative hepatic mRNA expression from 12-week-old WT^Cre+^, WT^Tg+^, and PGC-1α4^HepTg+^ male mice following 24 hrs of fasting (n=13). **(H)** Percentage of body weight loss in mice from A-F, calculated from values acquired before and after the 24-hr fast in each mouse. **(I)** Visceral adipose tissue weights. **(J)** mRNA expression levels in livers of 12-week-old male mice tail-vein injected with adeno-associated viral vectors (Control: AAV-shControl, PGC-1α4: AAV-shPGC-1α4) 2 week prior to a 24 hr fast. **(K)** Percentage of body weight gain, calculated from values acquired at baseline and 2 wks after expression of AAV-shControl or AAV-shPGC-1α4. **(L)** Visceral and (**M)** subcutaneous adipose tissue weights, **(N)** relative mRNA expression, **(O)** liver weight (%), **(P)** liver acylglycerol levels, **(Q)** serum total triglyceride levels, and **(R)** representative images of H&E stained liver sections in mice with reduced hepatic PGC-1α4 usinh shRNA delivered by AAV (or control virus) from **J**. Bars are mean ± SEM (n=9-11 mice per group). \**P* ≤ 0.05, \*\**P* ≤ 0.01, \*\*\**P* ≤ 0.001, \*\*\*\**P* ≤ 0.0001.

Prior to fasting, there were no significant differences in body weights between genotypes **(Supplementary Figure 8A-D**), yet it is noteworthy that transgenic PGC-1α4^HepTg+^ male mice lost more weight during the fast (**Figure 5I**). Consistently, acute *in vivo* reduction of hepatic PGC-1α4 (**Figure 5J**) led to weight gain (**Figure 5K**), and increased adipose tissue weights (**Figures 5L,M**), without any change on fat mass **(Supplementary Figure 9A-D**), and decreased *Fsp27* mRNA (**Figure 5N).** Paradoxically, knockdown of PGC-1α4 *in vivo* also increased liver weight, hepatic steatosis, and serum triglyceride (**Figure 5O-Q).** This unexpected steatosis may be the result of increased peripheral adipose tissue and/or what appears to be adaptive increases in hepatic *Cd36* and *Adrp* (**Supplementary Figure 9E,F**). Of note, the type of steatosis found in livers of shPGC-1α4 knockdown mice was primarily microsteatosis (**Figure 5R**), which is in line with the decrease in *Fsp27* expression (**Figure 5N**). These data illustrate that PGC-1α4 levels correlate well with hepatic macrosteatosis and *Fsp27* expression *in vivo*, but that any alteration in hepatic PGC-1α4 *in vivo* disrupts lipid metabolism.

### PGC-1α4 correlates with macrosteatosis in people living with MASLD and MASH

Since PGC-1α4 drove hepatic lipid accumulation and macrosteatosis in mice, we investigated whether PGC-1α4 expression is associated with steatosis or MASLD in humans. In general, levels of total *PGC-1α* mRNA transcripts are lower in MASLD^18–20^, but levels of individual isoforms have not been assessed. Using qPCR primers specific to Exon-1a (proximal promoter), or Exons - 1b and −1b’ (alternative promoter), we found increased transcription from the alternative promoter in liver samples from people living with MASLD and MASH (**Figure 6A**), correlating with increased expression of PGC-1α4 (**Figure 6B**). Non-quantitative PCR amplification of full length PGC-1α spliced transcripts showed that *PGC-1α-b* and *PGC-1α4* (alternative promoter) had low to no detection in people with low hepatic steatosis (0%, 0/7), but were inappropriately expressed in people with MASLD (43%, 6/14), and to some extent MASH (22%, 2/9) (**Figure 6C**), while incidence of *PGC-1α1* (proximal promoter) and *L-PGC-1α* (a liver specific isoform^28^) were consistent across all subjects.

**FIGURE 6.**
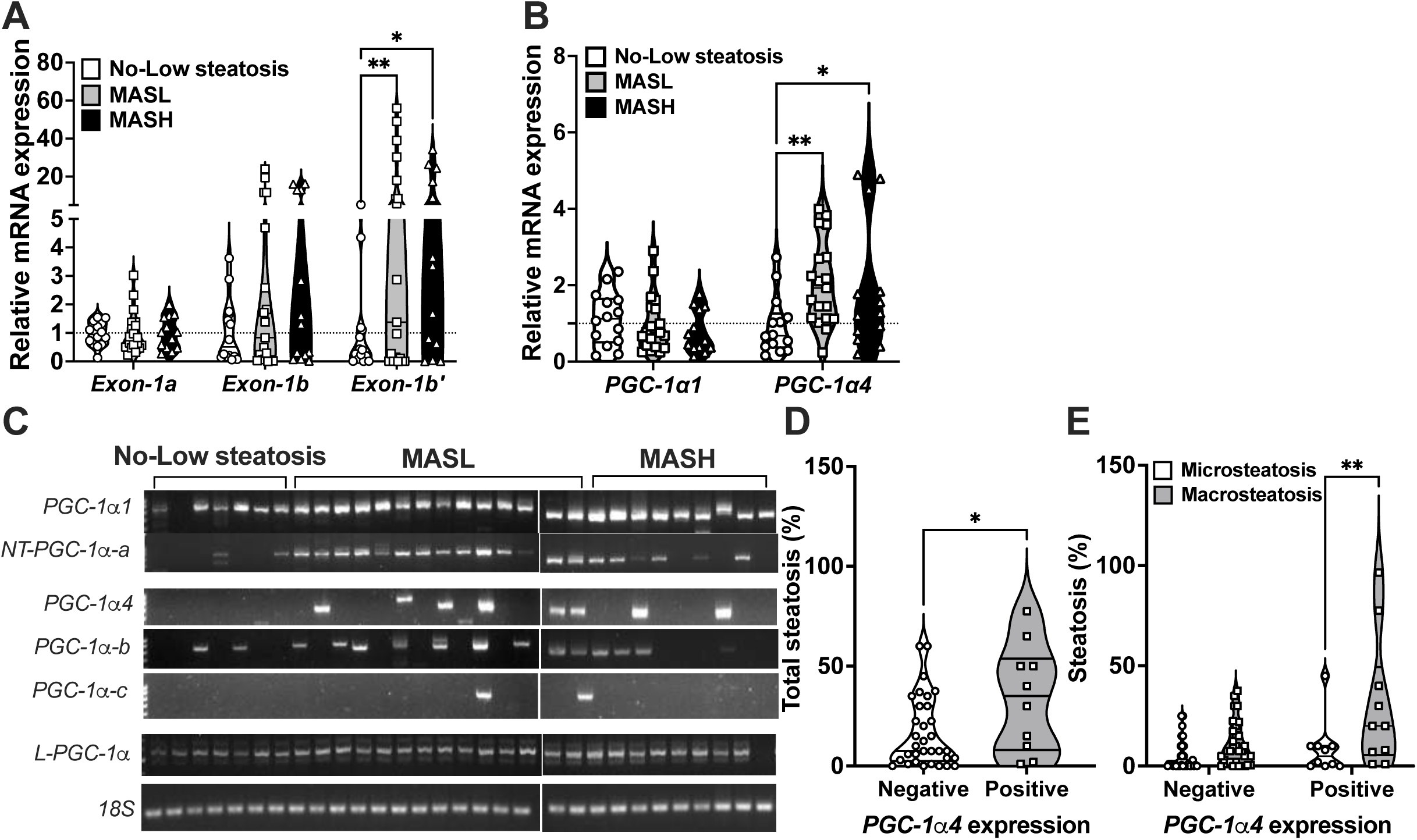
PGC-1α4 expression is higher in liver samples from patients with MASLD. **(A)** Relative mRNA levels of *PPARGC1A* transcripts containing exon-1a (proximal promoter), exon-1b (alternative promoter), or exon-1b′ (alternative promoter). **(B)** Relative mRNA expression of *PGC-1α1* and *PGC-1α4* isoforms. **(C)** Non-quantitative PCR of full length *PPARGC1A* transcripts in liver samples from patients with no/low steatosis (n=15, lanes 1-7), MAFL (n=20, lanes 8-21), and MASH (n=14, lanes 22-31). Percentage of liver steatosis **(D)** and micro- and macrosteatosis **(E)** in *PGC-1α4*-negative and *PGC-1α4*-positive liver samples across the three patient groups. Data are presented as violin plots with individual data points shown (min to max range). *P ≤ 0.05, **P ≤ 0.01, ***P ≤ 0.001, ****P ≤ 0.0001.

Since we discovered a mechanistic link between PGC-1α4 and hepatic macrosteatosis, we investigated whether there was a relationship between PGC-1α4 expression and level and type of steatosis in people with MASLD. We found that PGC-1α4 positive liver samples had both more steatosis (**Figure 6D**) and a higher prevalence of macrosteatosis (**Figure 6E**). Taken together, our data suggest that during MASLD progression, PGC-1α isoforms are differentially regulated and that inappropriate activation of the alternative *PPARGC1A* promoter leads to high PGC-1α4 which exacerbates hepatic macrosteatosis.

### High PGC-1α4 mimics a prolonged hepatic fasting response

As we also found mouse livers high in PGC-1α4 had reduced glycogen (**Figure 1I, Supplementary Figure 1E**), we thought that trapped lipid might direct hepatocytes toward using alternative energy sources. In line with this, levels of glucose, G6P, and glycogen were lower in PGC-1α4^HepTg+^ mice after refeeding (**Figure 7A-D,E,G)** and this was accompanied by increased expression of the glycogenolysis enzyme glycogen phosphorylase (*Pygl*) (**Figure 7F,H**), with no change in glycogen synthetase (*Gys2*). Glycolytic genes were also increased in livers of fasted PGC-1α4^HepTg+^ mice (**Figure 7I,K**). However, neither knockdown of hepatocyte PGC-1α4 *in vivo* **(Supplementary Figure 10A-C**) nor gain- or loss-of function *in vitro* **(Supplementary Figure 11A-P**) indicated a clear coactivator role for PGC-1α4 on glycolytic, gluconeogenic or glycogen metabolism genes

**FIGURE 7.**
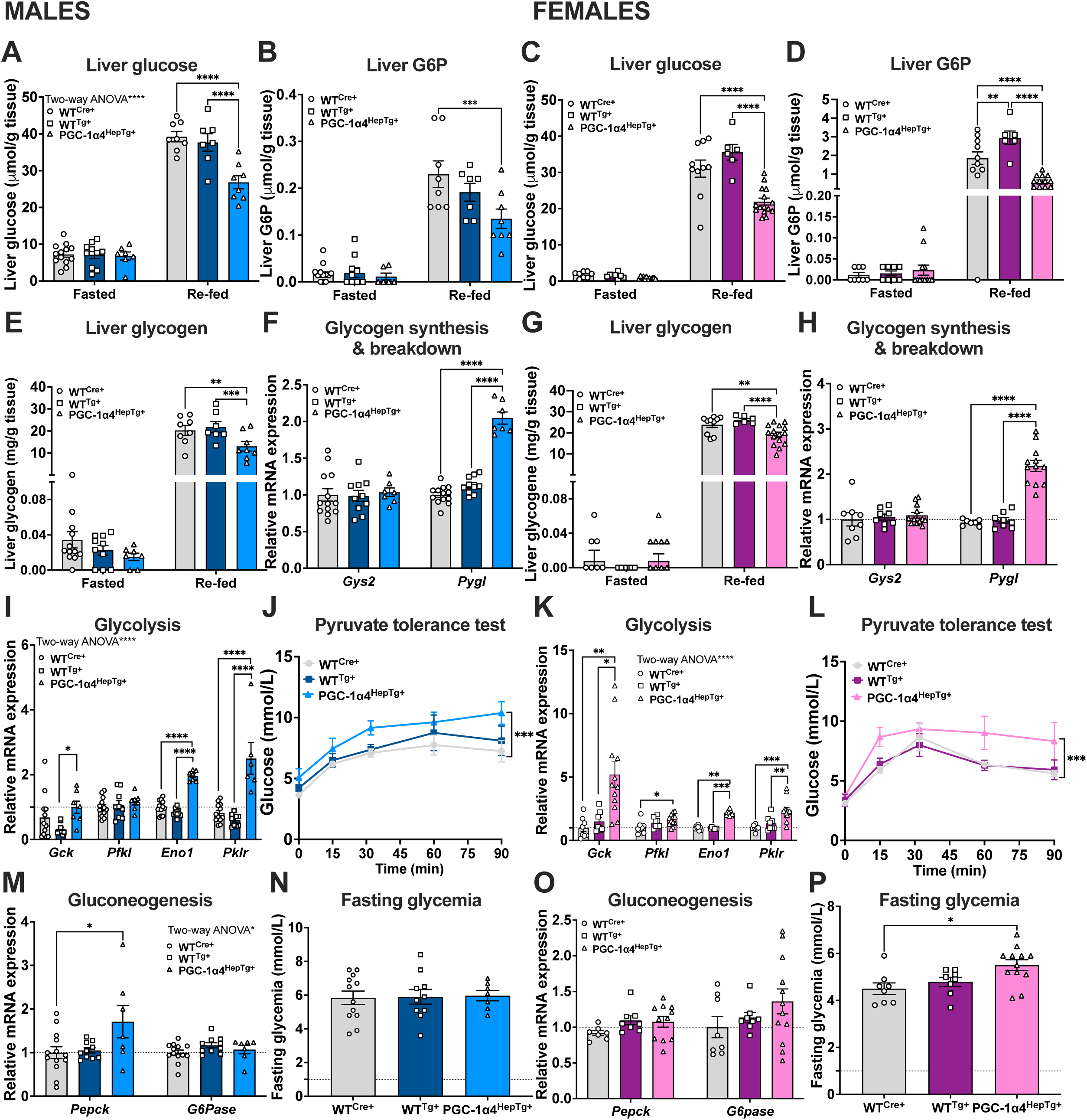
Sustained activation of PGC-1α4 in liver alters hepatic glucose homeostasis. **(A,C)** Liver glucose, (**B,D)** glucose-6-phosphate, and **(E,G)** glycogen levels in livers of 12-week-old WT^Cre+^, WT^Tg+^, or PGC-1α4^HepTg+^ male or female mice after 24 hours of fasting (Males, n=13; Females, n=7-12), or 6 hours re-feeding (Males, n=7-8; Females, n=6-15). **(F,H,I,K,M,)** Relative hepatic mRNA expression in fasted mice. **(J,L)** Pyruvate tolerance test in 32-week-old male or female mice (Males: n=4-5; Females: n=4-8). (N,P) Fasting glycemia 12-week-old WT^Cre+^, WT^Tg+^, or PGC-1α4^HepTg+^ male or female mice after 24 hours of fasting (Males, n=13; Females, n=7-12) Bars are mean ± SEM, \**P* ≤ 0.05, \*\**P* ≤ 0.01, \*\*\**P* ≤ 0.001, \*\*\*\**P* ≤ 0.0001.

Thus, we proposed that prolonged lipid trapping downstream of enhanced PPARy/*Fsp27* activity causes compensatory changes in liver to increase energy production from other sources. This explanation was consistent with the observed persistent fasting response, associated with increased glycolytic gene expression and blood glucose following pyruvate challenge in all PGC-1α4^HepTg+^ mice, and increased fasting glucose in females (**Figure 7I-P**). Overall, our combined *in vitro* and *in vivo* data support that the primary outcome of inappropriate hepatocyte PGC-1α4 expression is robust lipid trapping in large lipid droplets, which *in vivo* promotes an adaptive hepatic phenotype resembling a prolonged fasting state, further exacerbating steatosis.

### Increased hepatic PGC-1α4 combined with a western diet worsens macrosteatosis

It is well-known that prolonged liver steatosis correlates with liver inflammation, damage and fibrosis. 8-week-old PGC-1α4^HepTg+^ mice had steatosis with increased lobular inflammation and hepatocellular ballooning, but no fibrosis **(Supplementary Figure 12**). Considering our findings in humans, we sought to determine the impact of inappropriately high PGC-1α4 expression within a setting of MASLD. Interestingly, male and female PGC-1α4^HepTg+^ mice fed a diet high in saturated fat, fructose, and cholesterol gained less weight over time (**Figure 8A, Supplementary Figure 13A**) and had reduced body fat mass compared to controls (**Figure 8B, Supplementary Figure 13B**). Despite being leaner, PGC-1α4^HepTg+^ mice had larger livers (**Figure 8C, Supplementary Figure 13C**) and increased serum triglycerides (**Figure 8D, Supplementary Figure 13D**). *Pparγ* and *Fsp27* expression were increased in both sexes (**Figure 8E,F; Supplementary Figure 13E,F**), consistent with larger lipid droplets (mostly macrosteatosis) in livers PGC-1α4^HepTg+^ mice compared to controls (mostly microsteatosis) (**Figure 8G, Supplementary Figure 13G**). Notably, despite clear differences in steatosis type, total hepatic acylglycerols levels were similar across genotypes in males, and only significantly higher in female PGC-1α4^HepTg+^ mice (**Figure 8H, Supplementary Figure 13H**). There were no significant differences in serum aminoalanine transferase (ALT) (**Figure 8I), Supplementary Figure 13I**); however, genes associated with oxidative stress were higher in both sexes (**Figure 8J, Supplementary Figure 13J**, and male PGC-1α4^HepTg+^ livers also had increased inflammatory **(Supplementary Figure 14A**), immune cell **(Supplementary Figure 14B**), and fibrosis markers (**Supplementary Figure 14C**), but not females **(Supplementary Figure 14D-F**). These results indicate that excessive PGC-1α4 enhances lipid droplet size in the context of diet-induced MASLD, and this could be associated with an advanced transition toward MASH.

**FIGURE 8.**
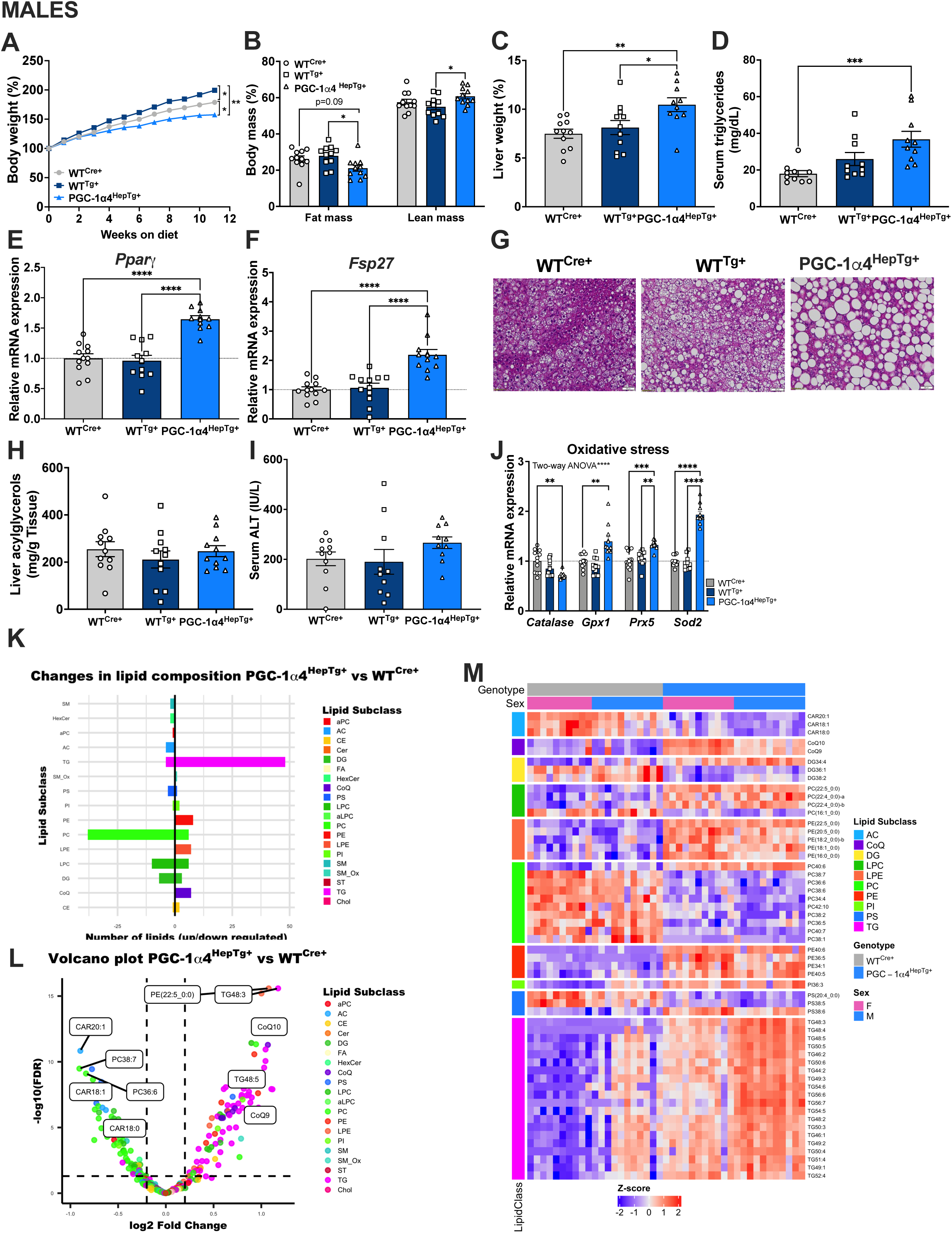
PGC-1α4 overexpression promotes macrosteatosis and oxidative stress in mice fed a Western Diet. **A)** Body weight, **(B)** Fat and lean mass measured by MRI, (C) Liver tissue weight, **(D)** serum triglycerides, **(E,F,J)** relative hepatic mRNA levels, **(G)** representative H&E stained liver sections, **(H)** liver acylglycerols, **(I)** serum ALT levels in male mice fed a high-fat, high-sucrose, high-cholesterol (Western) diet for 12 weeks. Data are expressed as mean ± SEM (n=11), *P ≤ 0.05, **P ≤ 0.01, ***P ≤ 0.001, ****P ≤ 0.0001. **(K)** main lipid class up and down regulated in the liver of PGC-1α4^HepTg+^ mice versus WT^Cre+^ mice; **(L)** volcano plot representing the changes among the 244 features annotated significatively altered in the liver of PGC-1α4^HepTg+^ mice versus WT^Cre+^ mice (adj.p.val < 0.05). Lipid species identified were: 1-alkyl,2-acylglycerophosphocholines= aPC, phosphocholines (sphingomyelins)= SM, Ceramide phosphocholines (sphingomyelins) (oxidized)= SM_Ox, Ceramides= Cer, Cholesterol and derivates= Chol, Diacylglycerols= DG, Diacylglycerophosphocholines= PC, Diacylglycerophosphoethanolamines= PE, Diacylglycerophosphoinositols= PI, Diacylglycerophosphoserines= PS, Fatty acids and conjugates= FA, Fatty acyl carnitines= AC, Monoacylglycerophosphocholines= LPC, Monoacylglycerophosphoethanolamines= LPE, Monoalkylglycerophosphocholines= aLPC, Simple Glc series= HexCer, Steryl esters= CE, Triacylglycerols= TG, Ubiquinones= CoQ. **(M)** heat maps of features changing significantly in the liver of PGC-1α4^HepTg+^ mice versus WT^Cre+^ mice (adj.p.val < 0.05). Each coloured cell represents the median value of a given feature across all samples in that group. Colour bars indicate magnitude of change in the level of that lipid, with red/blue depicting relatively higher/lower levels (within feature).

Since lipid type may influence the transition of MASL to MASH^53, 54^, we performed untargeted hepatic lipidomics to identify whether PGC-1α4 affects lipid class and/or composition. Principal Component Analysis (PCA) showed a reasonable separation between PGC-1α4^HepTg+^ and WT^Cre+^ mice, with no identified outliers **(Supplementary Figure 15A-B**). Linear regression revealed 396 significantly increased and 254 decreased features. Among these, 244 features were annotated as unique lipid species, and acyl chain composition was confirmed for 81 using MS/MS analysis. There was a marked increase in triacylglycerols (TAGs) in PGC-1α4^HepTg+^ livers, consistent with increased *Fsp27* expression and macrosteatosis. Interestingly, most of these TG features observed in the livers of PGC-1α4^HepTg+^ mice, is shown to be higher in humans with MASH, compared to MAFL^53, 54^, indicating prolonged expression of PGC-1α4 might be associated with more severe disease. The higher TAG was accompanied by increases in phosphatidylethanolamine (PE), while both phosphatidylcholine (PC) and phosphatidylserine (PS) were significantly decreased (**Figure 8K, L)**. This shift in membrane phospholipid composition reduces the PC:PE ratio, which drives membrane permeability, facilitating hepatocyte damage and impairing very-low-density lipoprotein (VLDL) secretion^55, 56^. Furthermore, we found elevated coenzyme Q (CoQ9/10) and reduced acylcarnitines in PGC-1α4^HepTg+^ livers (**Figure 8K, L)**. Coenzyme Q is decreases oxidative stress^57^ and its levels correlate with the increased oxidative stress response pathway. Reduced acylcarnitine suggests impaired long chain fatty acid transport into mitochondria and diminished lipid-oxidation^58^. Together with the *in vivo* liver phenotype, these results suggest that high PGC-1α4 expression not only drives neutral lipid accumulation, but also has profound changes on membrane phospholipid composition, which it might facilitate the transition of MASL to MASH.

## DISCUSSION

In this study, we show that PGC-1α4, a truncated variant of PGC-1α, is increased in fasting liver like canonical PGC-1α1, yet has a fundamentally distinct function. PGC-1α4 serves as a lipid-sparing signal, interacting with PPARγ to increase expression of lipid storage genes including *Fsp27* to promote lipid droplet fusion/expansion while reducing fatty acid oxidation and lipolysis. We propose that in a healthy fasting response, this regulatory mechanism ensures the availability of lipids stores that can be used for energy production after glycogen stores are depleted. However, dysregulated or prolonged expression of PGC-1α4 may lead to a persistent fasting-like metabolic state in liver. Sustained expression of PGC-1α4 is associated with the development of macrosteatosis, a hallmark of metabolic dysfunction. Correlations between elevated *PPARGC1A* alternative promoter activity, PGC-1α4 levels and macrosteatosis in individuals with MASLD suggest a direct link between aberrant PGC-1α4 expression and the worsening of liver disease.

In contrast to canonical PGC-1α1, PGC-1α4 did not induce lipid oxidation-related gene transcription, oxidative metabolism, or mitochondrial biogenesis in primary hepatocytes. This is in line with studies in mouse myocytes, where it does not alter expression of established canonical PGC-1α1 target genes^30, 31, 59^. Despite sharing some structural domains with canonical PGC-1α1, PGC-1α4 is a truncated version that has a different exon 1 and lacks the majority of the C-terminus important for nuclear localization, binding to the mediator complex, and its role in RNA splicing, potentially explaining reduced transcriptional activity for PGC-1α4^27^. However, this could also be due to its predominant cytoplasmic localization^16^. We show evidence here that nuclear translocation of PGC-1α4 can increase interactions with nuclear co-factors, including PPARγ, facilitating its role as a transcriptional coactivator.

Our findings appear to diverge from the canonical view of PPARγ as an anabolic signal engaged primarily under nutrient-rich conditions such as feeding. The paradox surrounding PPARγ largely reflects its extensive characterization in adipose tissue. In this context, nutrient availability or pathological feeding increases PPARγ expression, while fasting reduces it, a pattern that mirrors adipocyte *Fsp27*^4, 60, 61^. In liver, although most studies (including ours) report a decrease in PPARγ mRNA in fasting mice^4–6^, we show here that hepatic PPARγ protein levels increase during fasting. Consistently, *Fsp27* expression is also elevated in fasting liver^4, 5, 48^. PPARγ is generally required for hepatic lipid accumulation under many metabolic contexts^62^; yet, its role in fasting-induced steatosis has been challenged^4^. However, in this context, CREB might compensate^48^. Nevertheless, our *in vitro* experiments reveal that in response to various stimuli (glucagon, rosiglitazone, free fatty acids), *Fsp27* expression is dependent on PPARγ. We propose that the coordinated induction of PGC-1α proteins and PPARγ helps to manage the dual anabolic–catabolic nature of hepatic fasting, where coactivation of PPARγ drives a gene program that facilitates fatty acid uptake and triglyceride storage, promoting a reservoir for fatty acids mobilized from adipose lipolysis used for energy during prolonged fasting.

PGC-1α4’s promotion of hepatic lipid storage is in direct opposition to the catabolic function of canonical PGC-1α1. Thus, it was surprising to find that glucagon and fasting simultaneously induce mRNA expression of both isoforms. However, we observed that PGC-1α4 protein has delayed expression and prolonged stability in hepatocytes, in contrast to the short-lived and cyclic expression of PGC-1α1 protein. Combining knowledge of the known and recently discovered functions for these two isoforms, we propose that PGC-1α1 is rapidly induced at the onset of fasting, playing an acute role in stimulating lipid uptake, but also nutrient catabolism, oxidative metabolism, mitochondrial biogenesis, and gluconeogenesis, yet its expression quickly declines. In contrast, the delayed and extended PGC-1α4 expression during extended fasting can promote lipid sequestration and limit its catabolism during periods when glucose/glycogen are still an available energy resource. The reappearance of PGC-1α1 after prolonged fasting may help to later shift the balance toward increased catabolism of the stored lipid fuels. Thus, coordinated action of these two isoforms might be particularly important during extended fasting periods, where long-term conservation of fat stores is critical. Differential PGC-1α1 and PGC-1α4 expression and function ensure a healthy balance between energy mobilization and conservation.

We also show that dysregulated expression of PGC-1α4 leads to steatosis. Hepatocytes overexpressing PGC-1α4 cannot efficiently release stored acylglycerols or oxidize fatty acids, and we attribute these deficiencies to the trapping of lipids within droplets. Interestingly, hepatocyte-specific PGC-1α4 transgenic mice also have reduced body weight, leading us to speculate that expression of PGC-1α4 promotes a ‘permanent fasting’ state in liver. This could be a signal for increased adipose tissue lipolysis, leading to increased fatty acid uptake by liver, further exacerbating hepatic steatosis. Surprisingly, reducing hepatic PGC-1α4 expression also led to lipid accumulation in liver. If reduced PGC-1α4 expression consequently induces a ‘fed’-like state, this is consistent with observed increases in hepatic acylglycerols and weight gain. When combined with a Western diet, increased hepatic PGC-1α4 exacerbates lipid droplet expansion, a hallmark of macrosteatosis, representing 80-90% of the steatosis seen in MASLD^63–65^. *Fsp27* expression is also positively associated with the severity of MASLD in mice and in humans^66^. We found that people living with MASLD who also have high hepatic PGC-1α4 expression have increased steatosis with more prevalent macrosteatosis. Thus, while the PGC-1α4/PPARγ axis may be essential for efficient hepatic lipid handling, within a pathological context this mechanism may become maladaptive, whereby sustained activation promotes excessive lipid accumulation and exacerbates metabolic dysfunction. This is consistent with our *in vivo* data showing that increased lipid storage caused by high PGC-1α4 can eventually lead to oxidative and inflammatory damage, which may facilitate the transition from MASLD to MASH.

Our lipidomics analysis supports our model by revealing a pronounced shift toward triglyceride storage when hepatic PGC-1α4 is high. Notably, triglyceride accumulation is closely associated with the transition from MASL to MASH^53, 54^. Mechanistic studies suggest that impaired phosphatidylcholine (PC) synthesis during triglyceride accumulation reduces the surface area-to-volume ratio of lipid droplets, favoring macrosteatosis^67^. In line with this, reduced hepatic PC:PE ratio has been consistently linked to altered membrane dynamics, hepatic steatosis, and progression to MASH in the context of MASLD^55^. Beyond phospholipid remodeling, our data point to increased oxidative stress and overall decreased fatty acid oxidation, which may reflect adaptive responses to mitochondrial dysfunction or redox imbalance. Together, these findings indicate that PGC-1α4-driven lipid changes converge on membrane phospholipid composition and triglyceride metabolism, potentially destabilizing hepatocellular membrane integrity as well as promoting lipid accumulation, further contributing to disease progression.

## Conclusion

Our findings highlight the dual nature of PGC-1α proteins in liver metabolism: essential for maintaining energy balance during fasting, but potentially harmful when dysregulated in pathological contexts. Under healthy conditions, PGC-1α4/PPARγ facilitates storage of fat in liver during prolonged fasting, supporting continued energy metabolism in times of nutrient scarcity. However, unregulated expression impairs the intricate balance of catabolism and storage. Increased interactions between PGC-1α4/PPARγ, exacerbated by increased glucagonemia and circulating lipids associated with metabolic disease, could play a critical role in the progression from simple steatosis to more advanced liver disease, such as MASH. Our study shows that exploring the complex mechanisms that control fasting can offer a better understanding of the causes metabolic liver diseases including MASLD.

## Supporting information

Graphical Abstract

