## Supplementary figures and images for "Glucagon-induced PGC-1α4/PPARγ promotes hepatic lipid storage and macrosteatosis in fasting and MASLD"

### Graphical Abstract

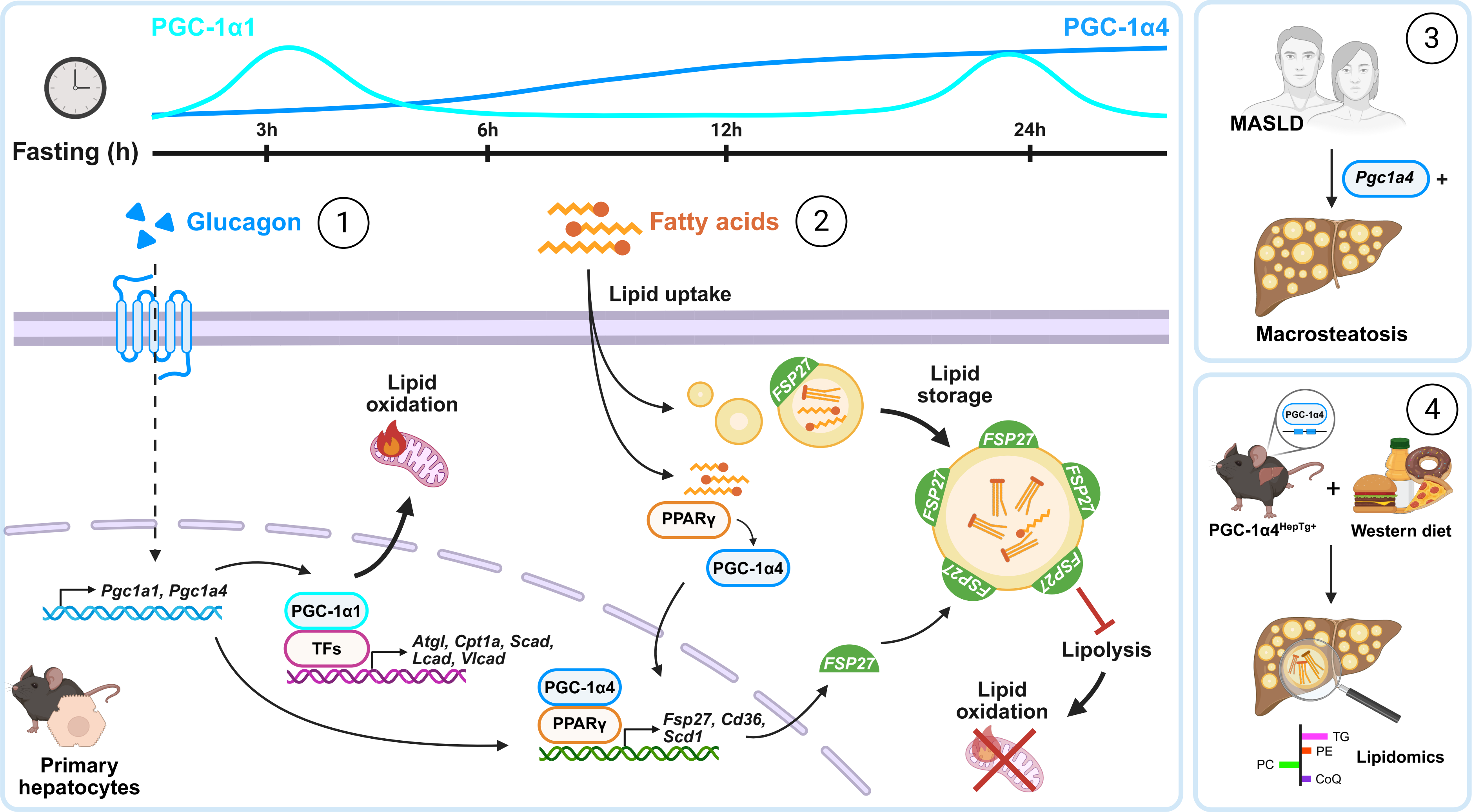
