## Supplementary figures, legends, tables for "Glucagon-induced PGC-1α4/PPARγ promotes hepatic lipid storage and macrosteatosis in fasting and MASLD"

#### SUPPLEMENTARY FIGURES, TABLES AND LEGENDS

### SUPPLEMENTARY FIGURE 1

#### FEMALES

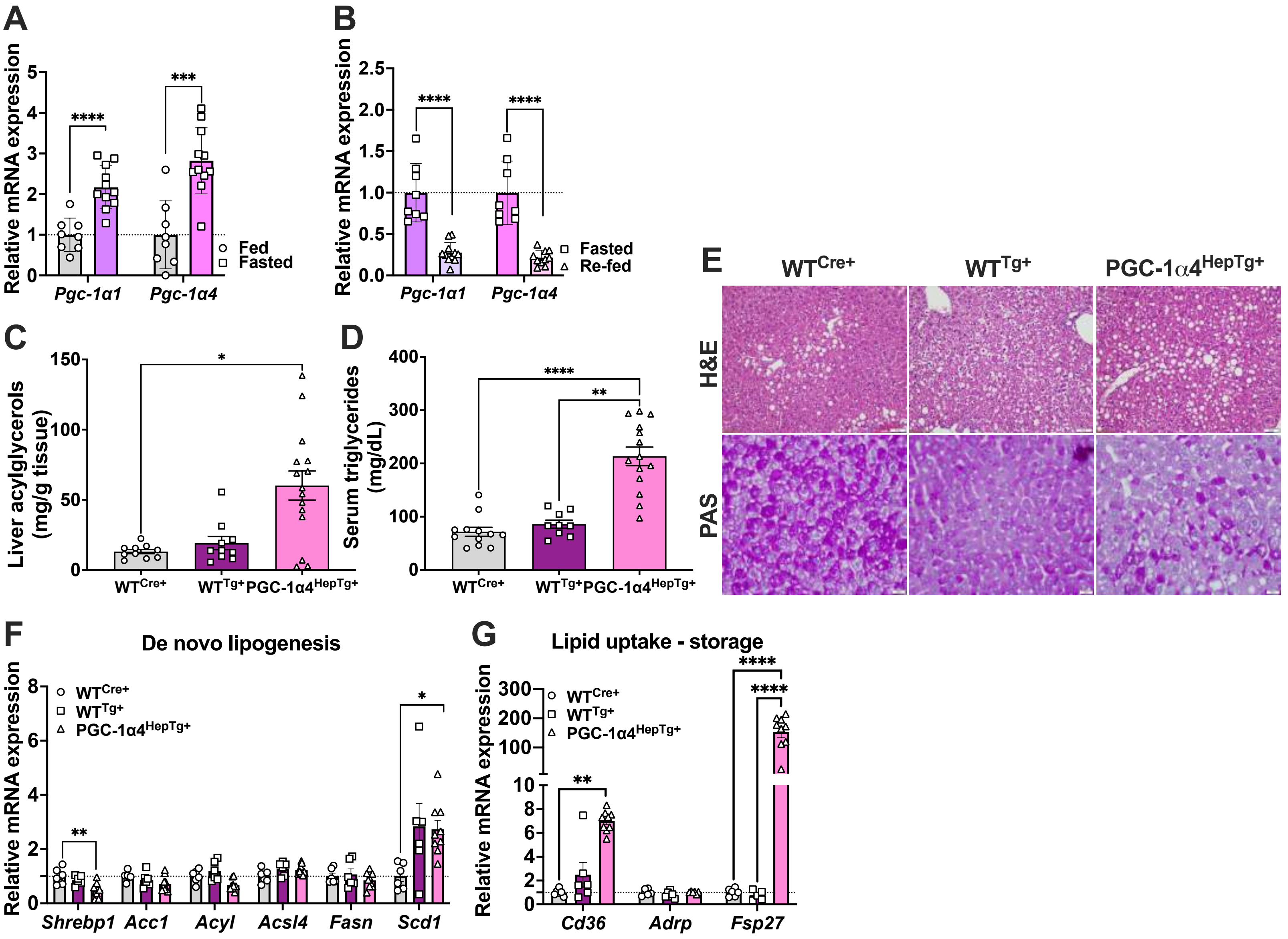

### SUPPLEMENTARY FIGURE 2

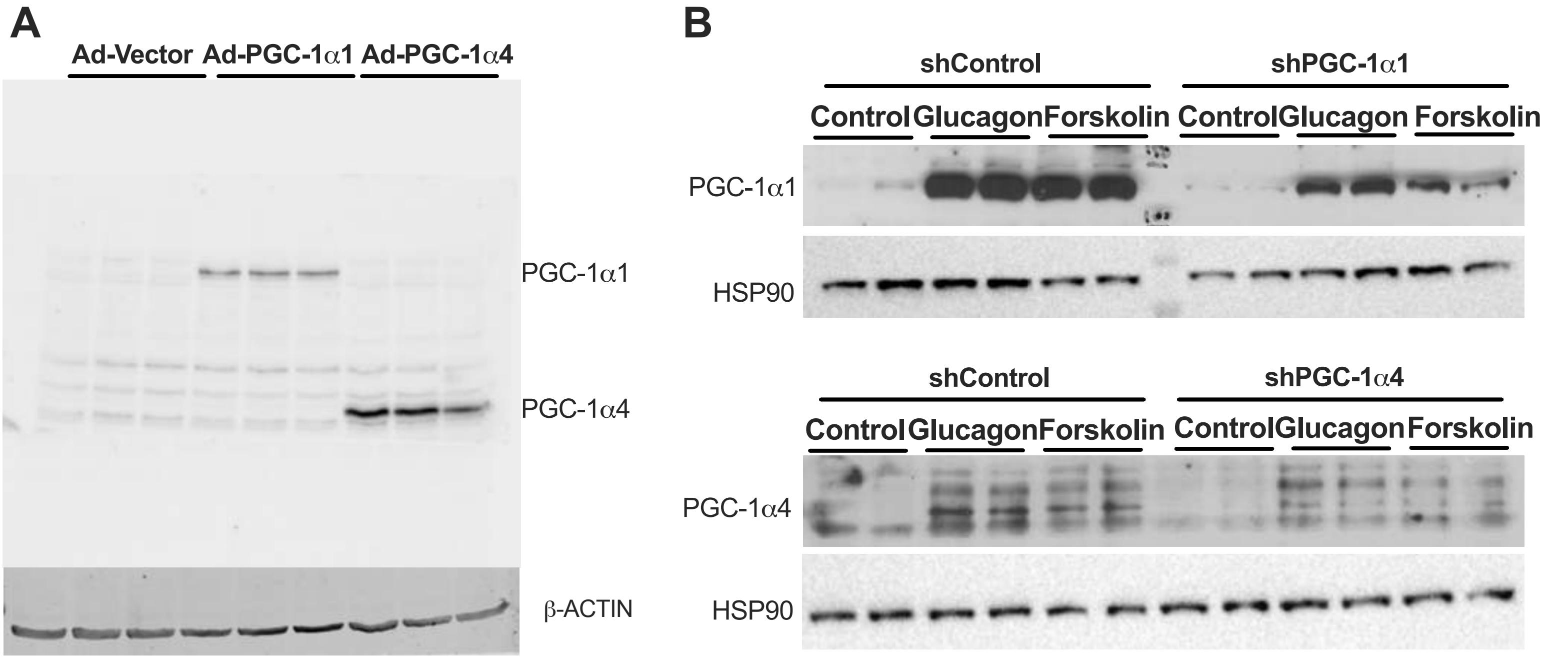

### SUPPLEMENTARY FIGURE 3

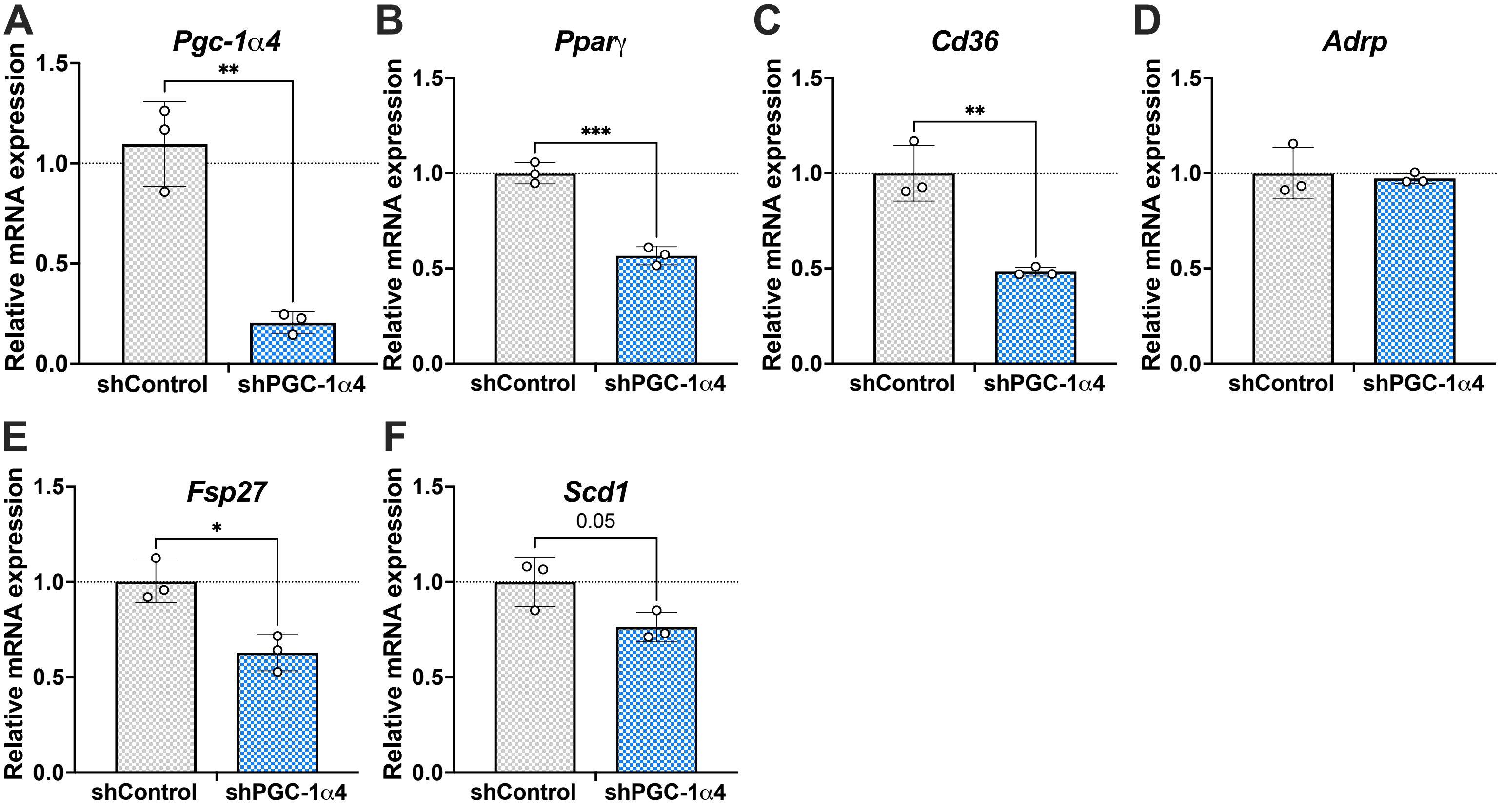

### SUPPLEMENTARY FIGURE 4

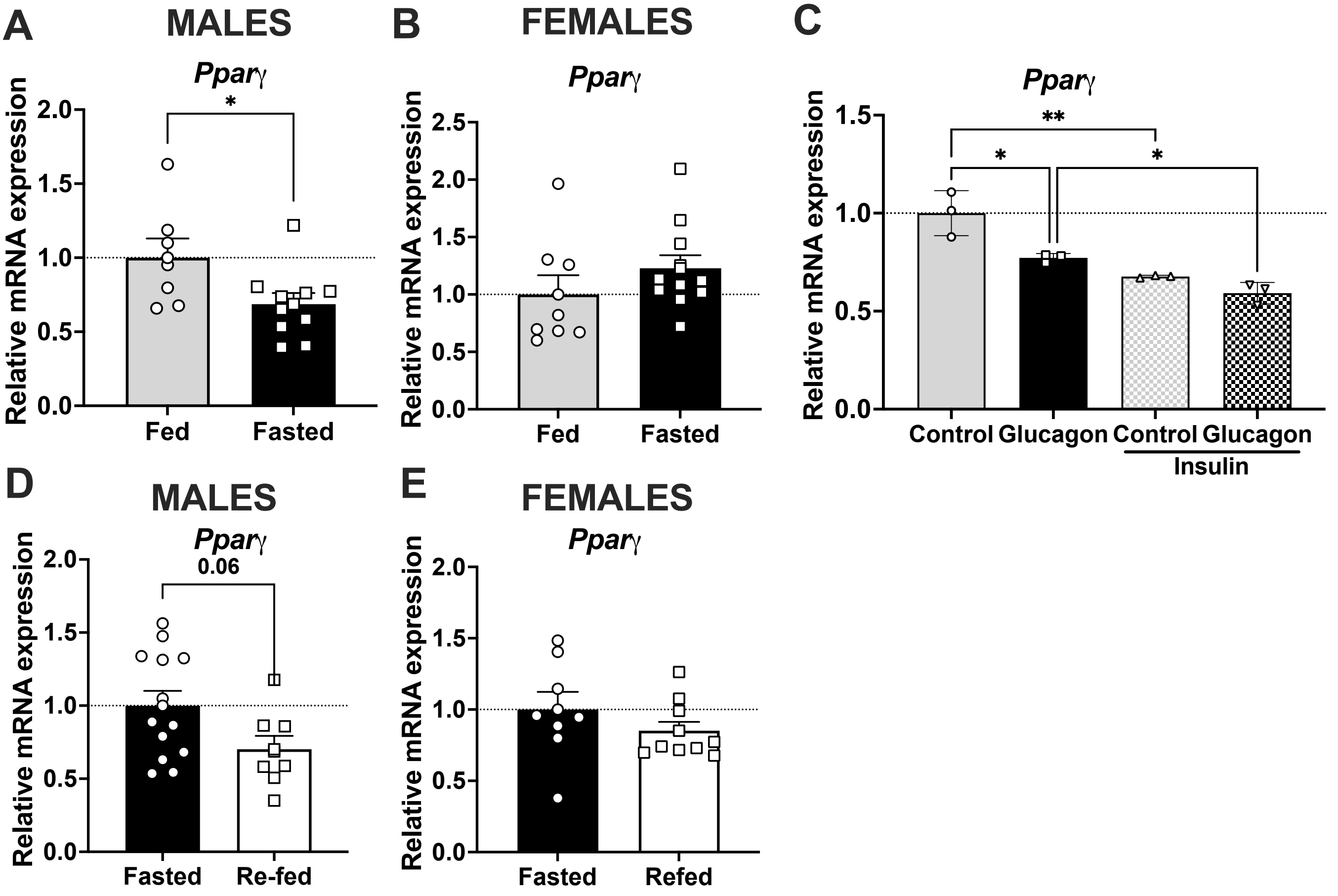

### SUPPLEMENTARY FIGURE 5

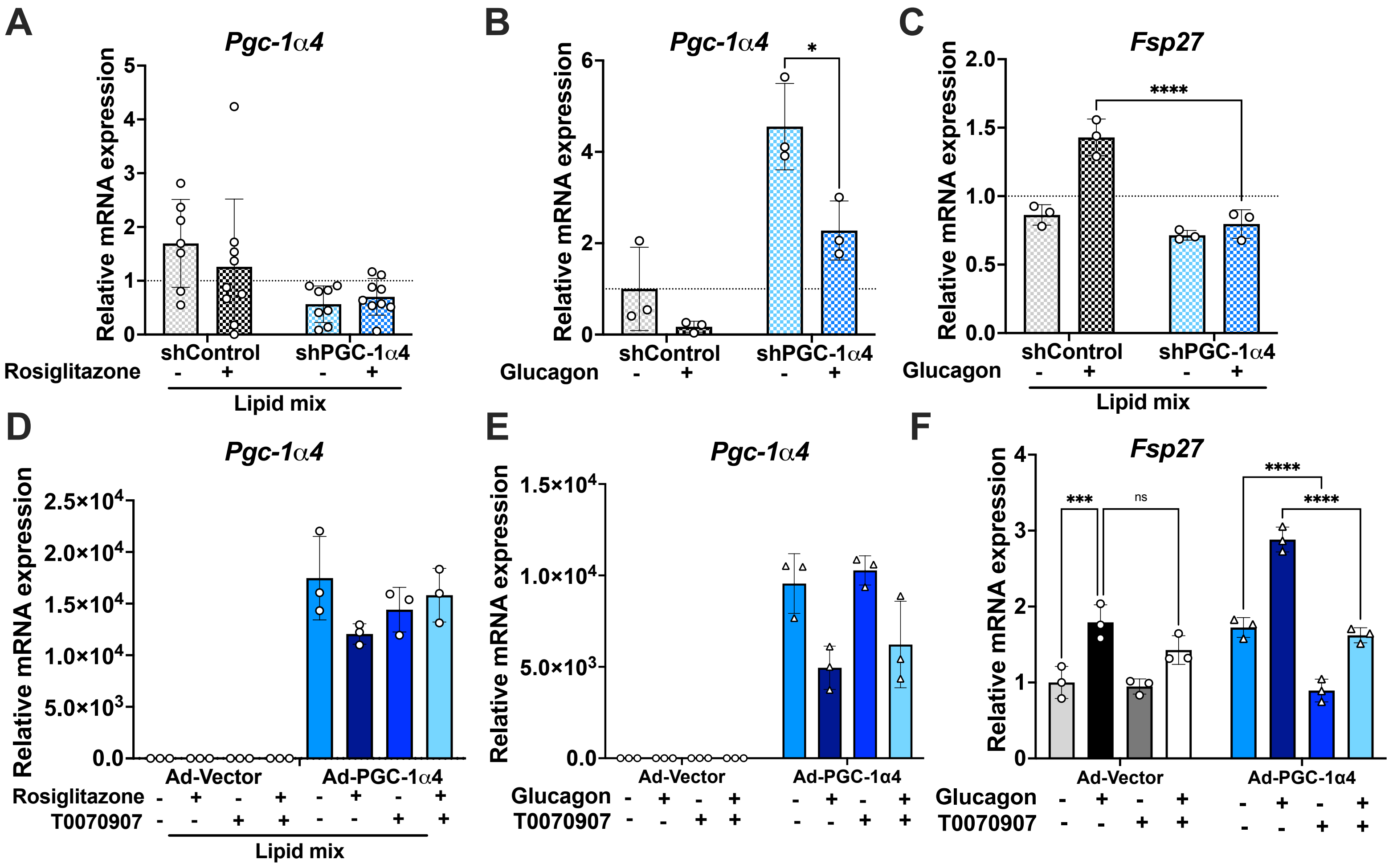

### SUPPLEMENTARY FIGURE 6

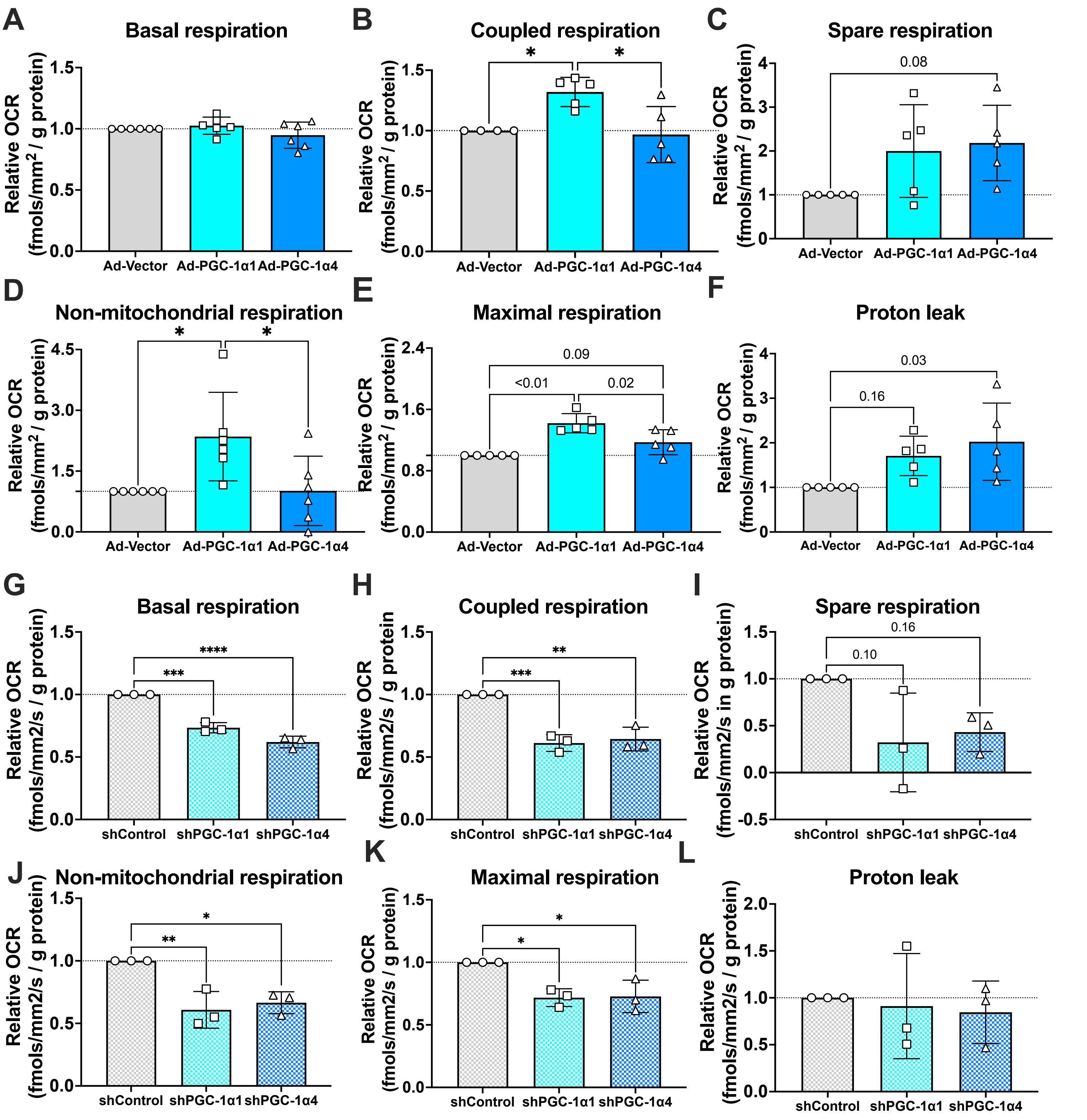

### SUPPLEMENTARY FIGURE 7

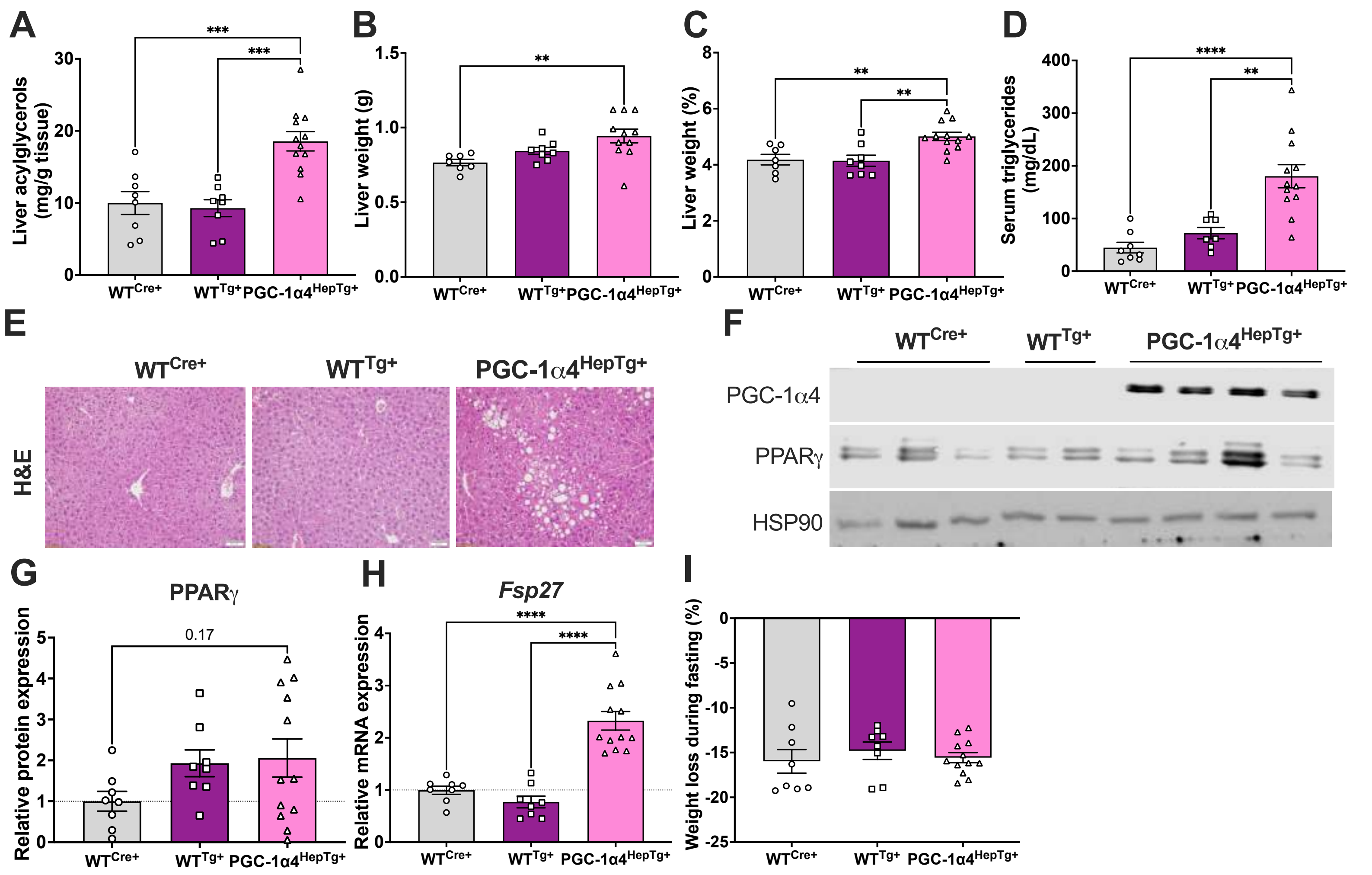

### SUPPLEMENTARY FIGURE 8

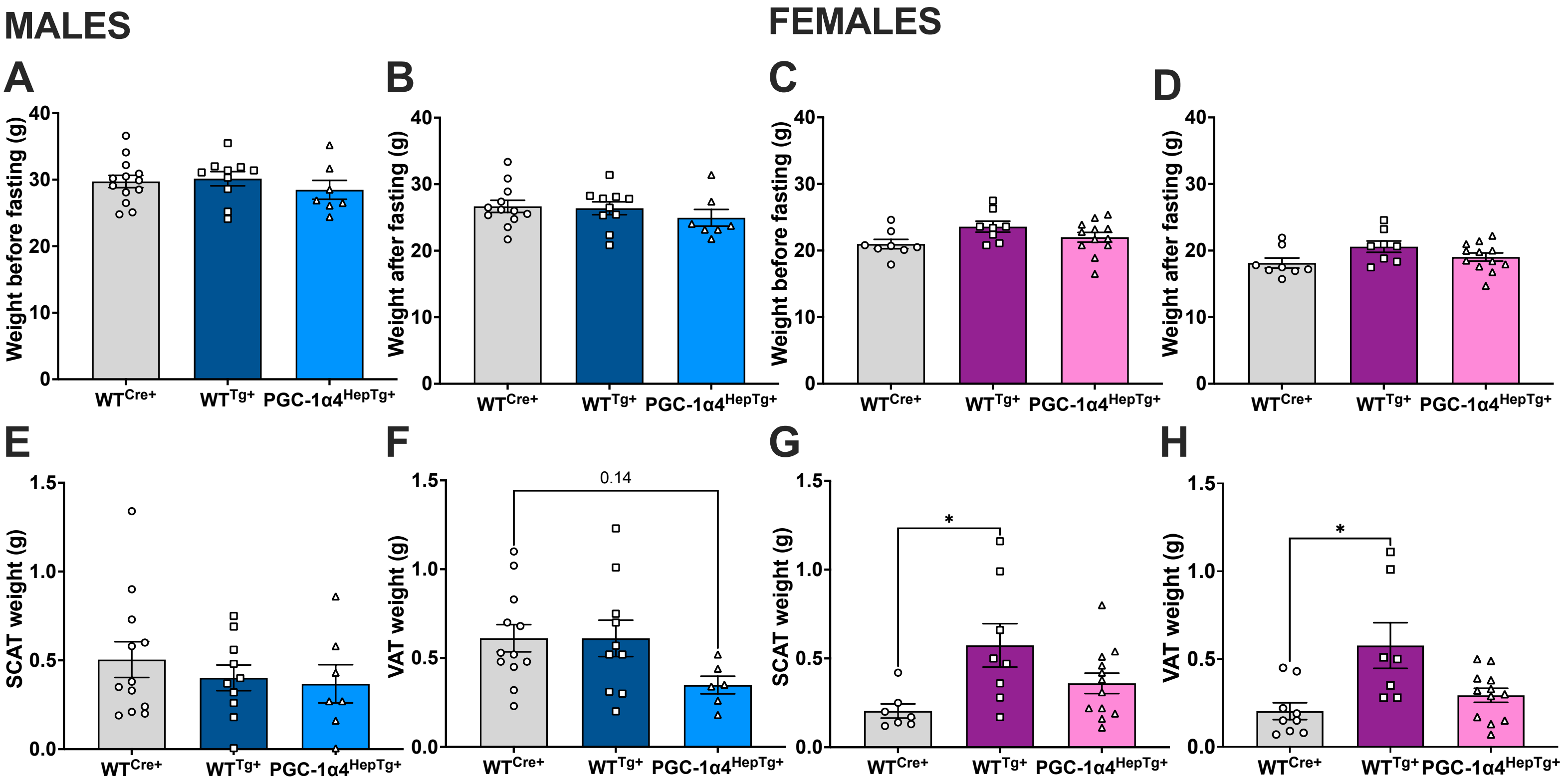

### SUPPLEMENTARY FIGURE 9

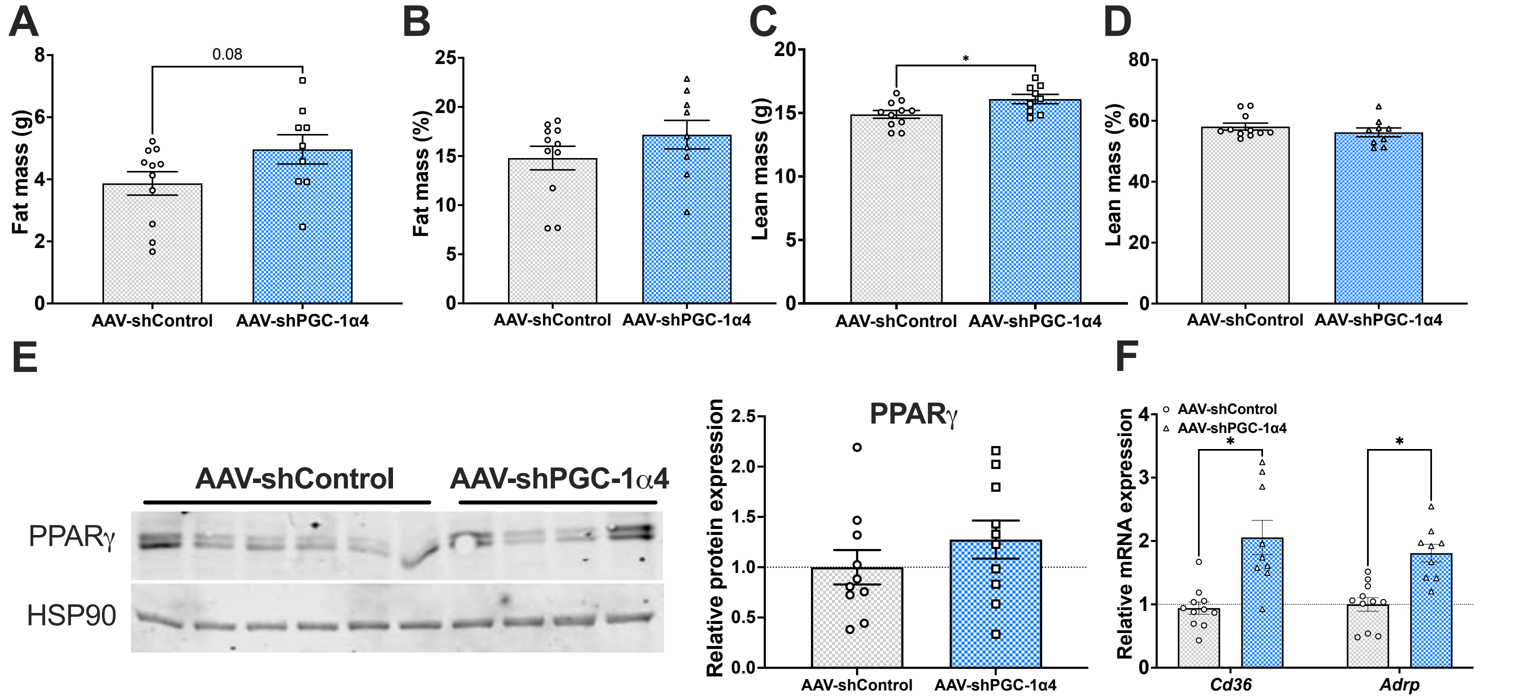

### SUPPLEMENTARY FIGURE 10

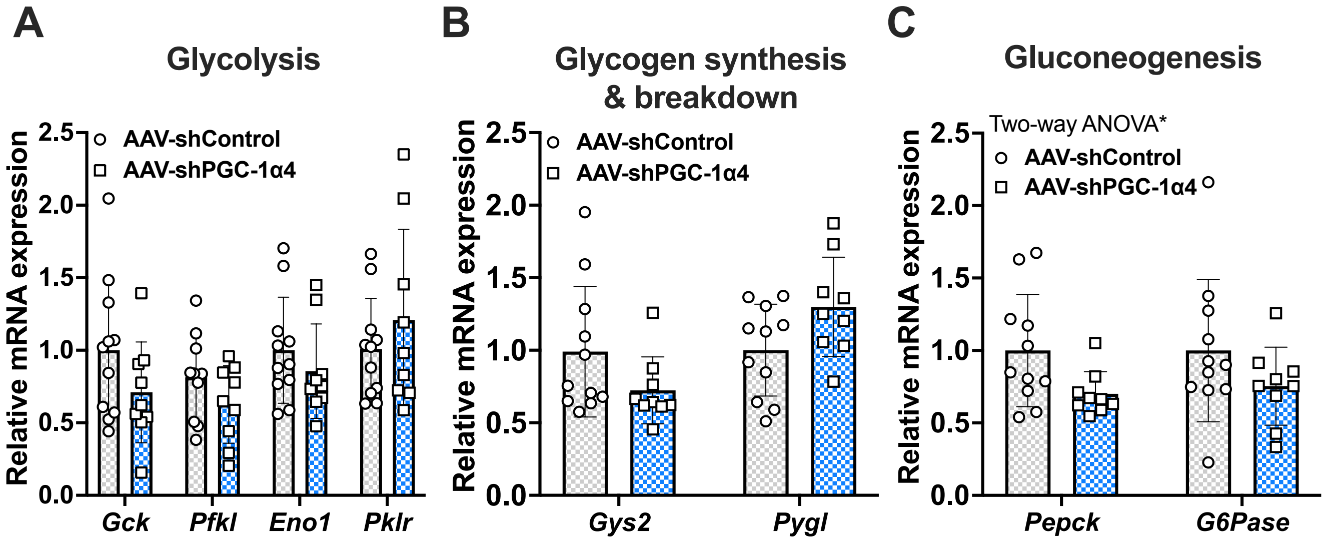

### SUPPLEMENTARY FIGURE 11

#### Glycolysis

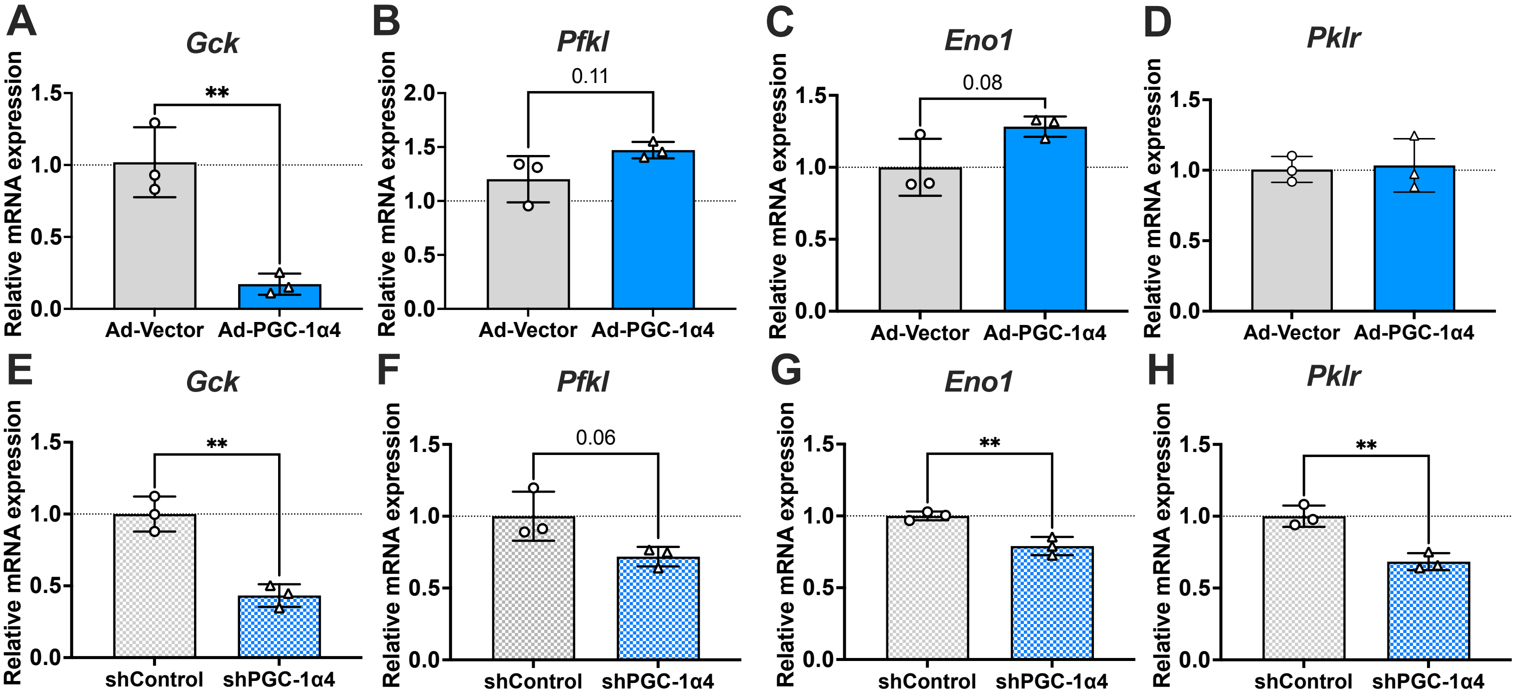

#### Glycogen syntesis & breakdown

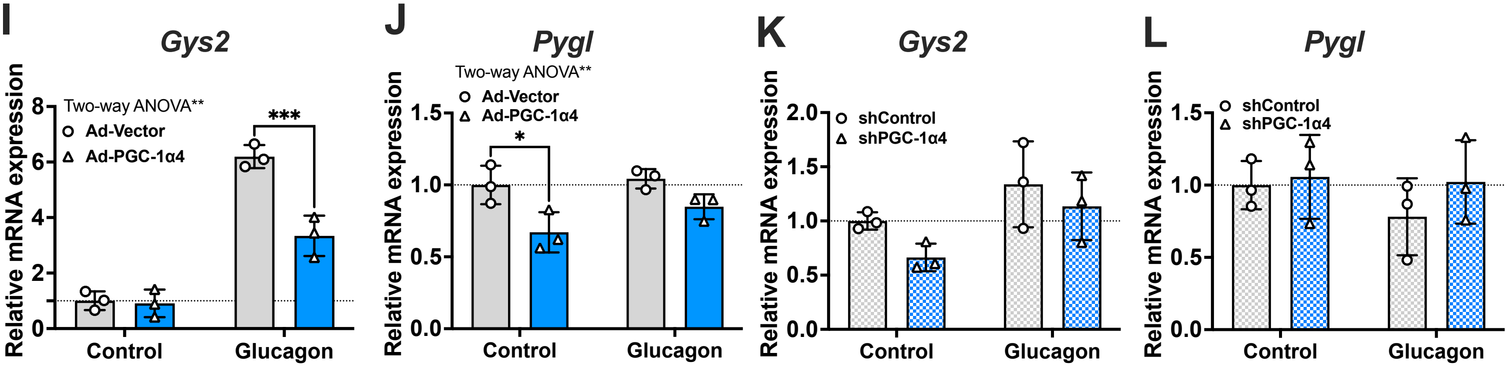

#### Gluconeogenesis

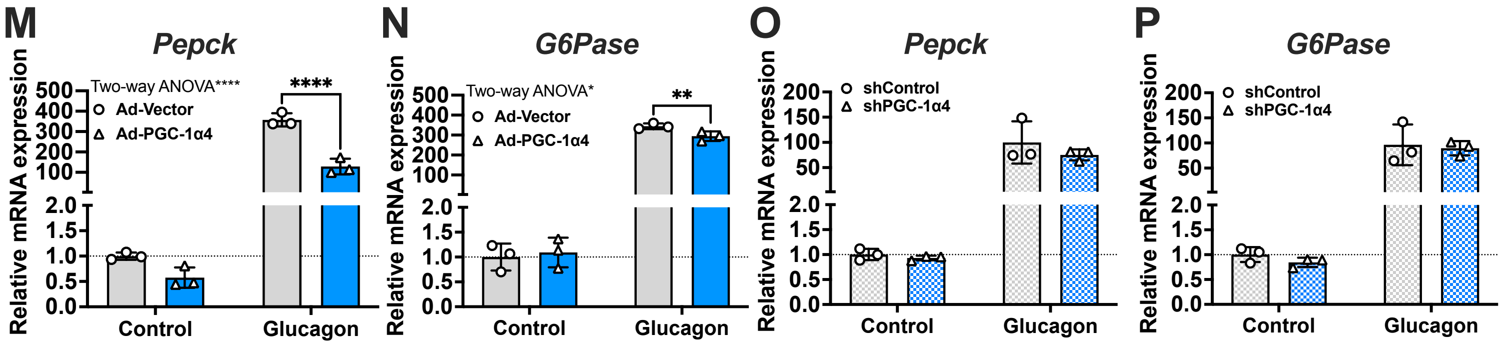

### SUPPLEMENTARY FIGURE 12

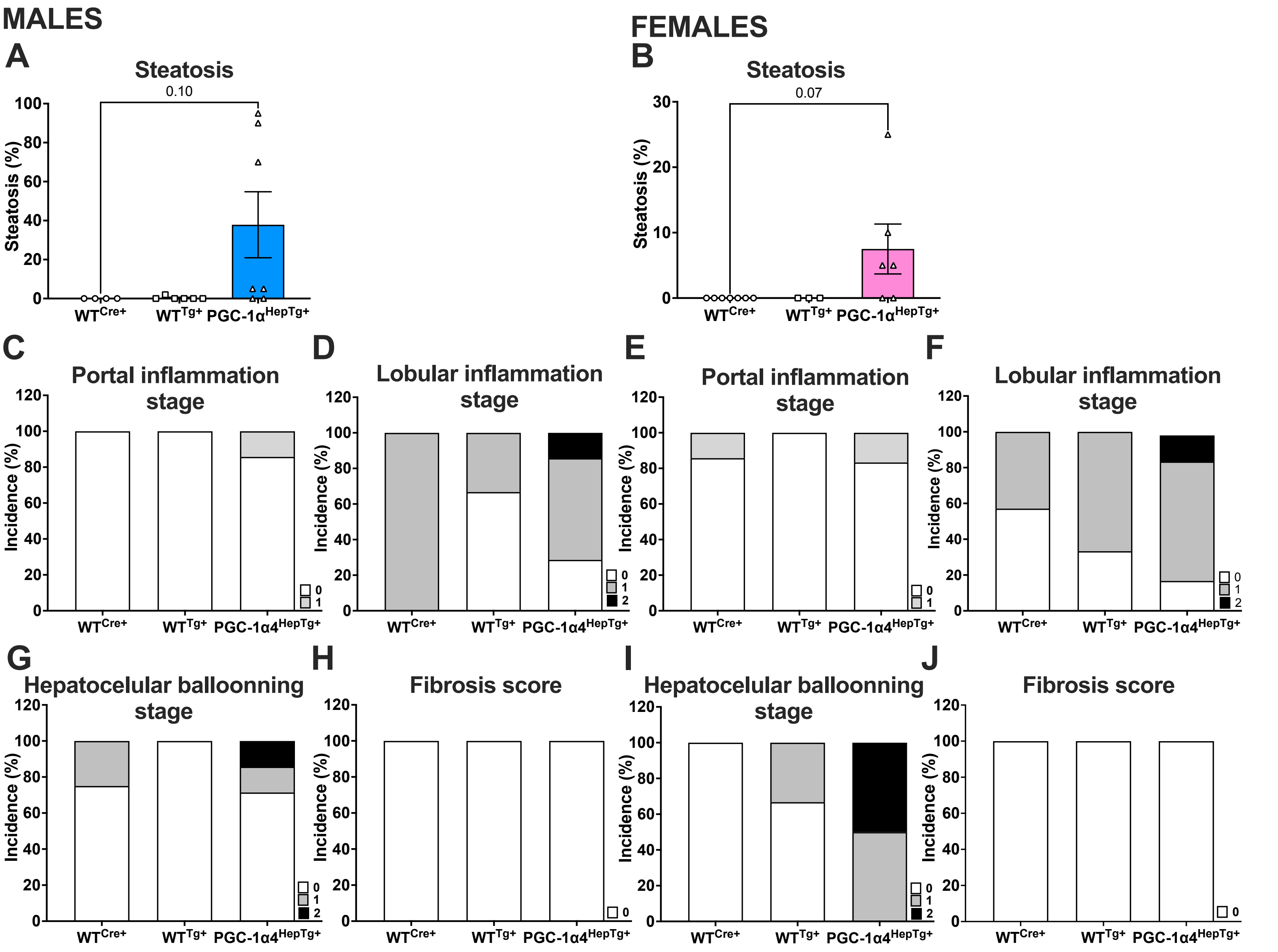

### SUPPLEMENTARY FIGURE 13

#### FEMALES

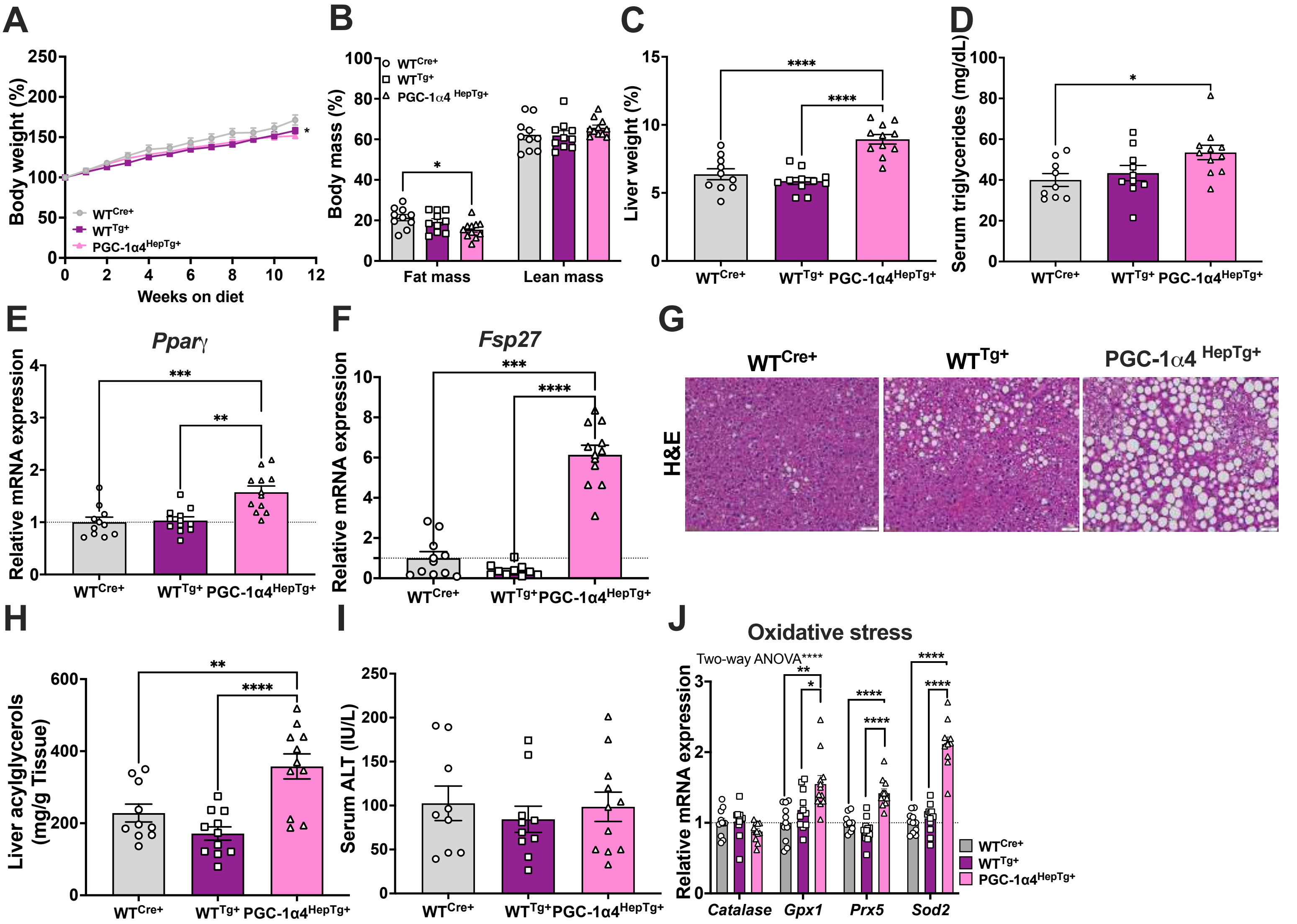

### SUPPLEMENTARY FIGURE 14

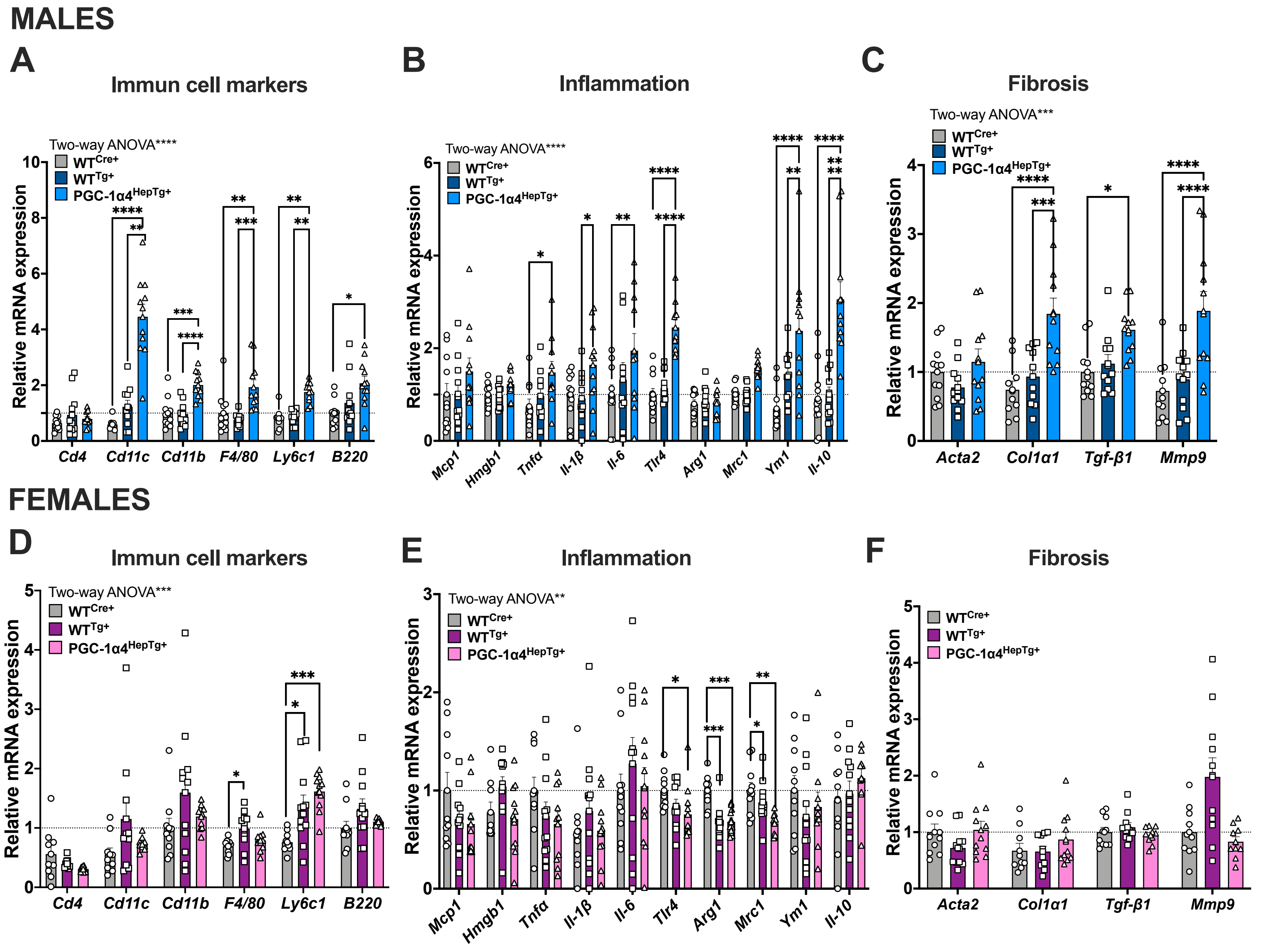

### SUPPLEMENTARY FIGURE 15

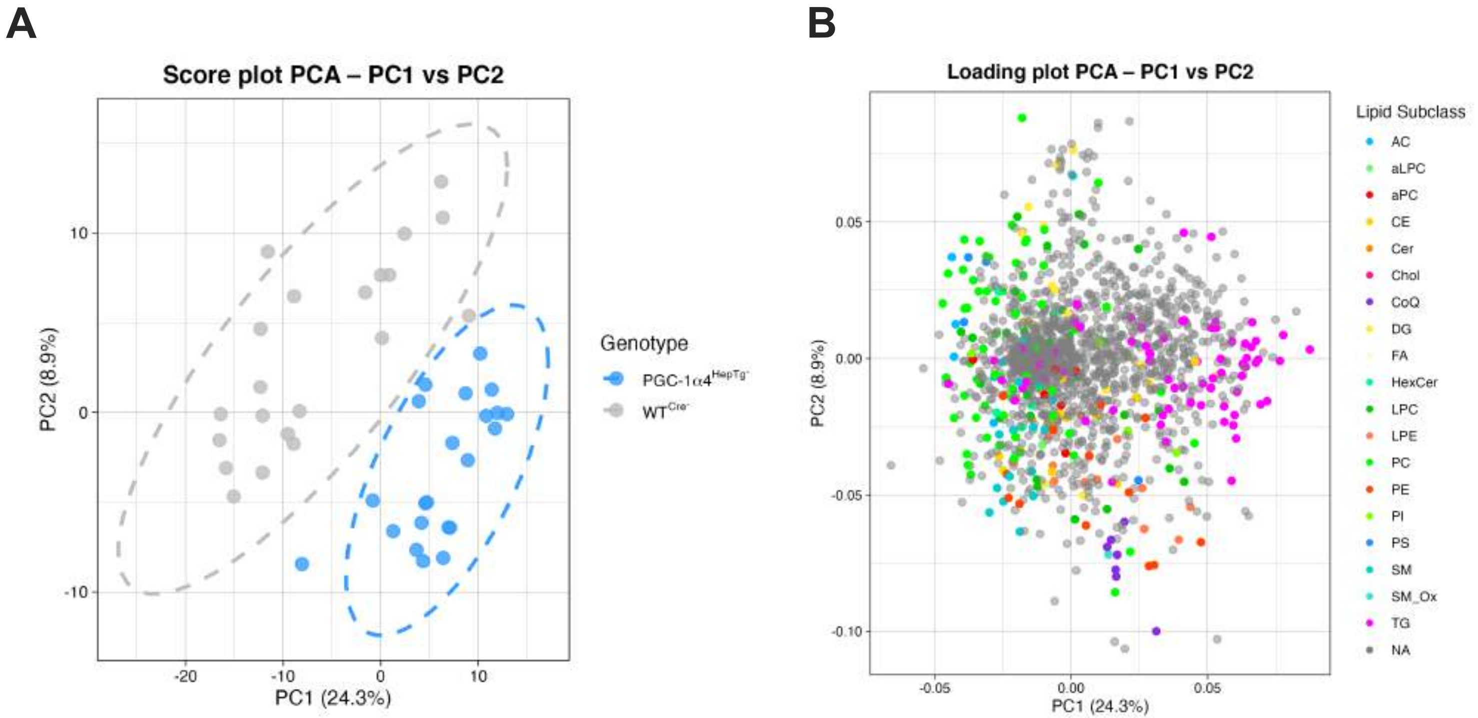

**SUPPLEMENTARY FIGURE 1. PGC-1α1** ***and*** **PGC-1α4** ***are induced by*** **fasting and repressed by feeding in mouse liver.**

Relative hepatic mRNA levels in the livers of 12-week-old C57BL/6J male mice **A)** fasted for 24 hrs (n=8-11) or **B)** fasted for 16 hrs and re-fed for 6 hrs (n=8-11). **(C)** Total liver acylglycerols, **(D)** serum total triglycerides, and **(E)** representative H&E and PAS staining of liver sections from 8-12 weeks old male mice (n=6-15). **(F,G)** Relative mRNA expression in the livers of 12-week-old male mice (n=6-9). Bars represent means ± SEM, $*P \leq 0.05$, $**P \leq 0.01$, $***P \leq 0.001$, $****P \leq 0.0001$.

**SUPPLEMENTARY FIGURE 2.** ***Validation of PGC-1α1 and PGC-1α4 overexpression and silencing in primary hepatocytes in-vitro.***

**(A)** Protein expression of *PGC-1α1* (~110 kDa) and *PGC-1α4* (~38 kDa) in primary hepatocytes infected with adenoviral vectors (Ad-Vector, Ad-PGC-1α1, and Ad-PGC-1α4) for 48 hours. **(B)** Protein expression following silencing of *PGC-1α1* and *PGC-1α4* by adenoviral vectors (shControl, shPGC-1α1 and shPGC-1α4) for 72 hours by western blotting.

**SUPPLEMENTARY FIGURE 3.** ***The effect of reducing PGC-1α4 expression on PPARγ and its target genes.***

(A-F) Relative mRNA expressions in PGC-1α4 silenced primary hepatocytes for 72 hrs. Bars represent the mean ± SD of four to ten biological replicate. $*P \leq 0.05$, $**P \leq 0.01$, $***P \leq 0.001$, and $****P \leq 0.0001$.

**SUPPLEMENTARY FIGURE 4.** ***The analysis of hepatic Pparγ mRNAexpression in fasting***

Relative mRNA expression of *Pparγ* in livers of 12-week-old C57BL/6J **(A)** male and **(B)** female mice after 24 hrs of fasting. Data are expressed as means ± SEM (n=7-12). **(C)** Relative *Pparγ* mRNA levels in primary hepatocytes treated with 50 nM glucagon for 24 hrs +/- 100 nM insulin for the last 12 hrs. Data are mean ± SD representative of 2-3 independent experiments done in triplicate. Relative *Pparγ* mRNA levels in livers of **(D)** male and **(E)** female C57BL/6J ($WT^{Cre+}$) mice re-fed for 6 hrs following a 16-hr fast. Data are expressed as means ± SEM (n=7-12). $*P \leq 0.05$, $**P \leq 0.01$, $***P \leq 0.001$, and $****P \leq 0.0001$.

**SUPPLEMENTARY FIGURE 5.** ***The effect of modulating PGC-1α1 and PGC-1α4 expression on Fsp27 mRNA expression in primary hepatocytes in response to fasting signals.***

**(A)** Relative mRNA levels in in primary hepatocytes expressing shPGC-1α4 or shControl treated with rosiglitazone and the lipid mix (n=2). (B) Relative mRNA levels in in primary hepatocytes expressing shPGC-1α4 or shControl treated with glucagon (n=1). (C) Relative mRNA levels in in primary hepatocytes expressing shPGC-1α4 or shControl treated with glucagon and the lipid mix (n=1). **(D)** Relative mRNA expression levels in primary hepatocytes expressing Ad-**PGC-1α4** or Ad-Vector control pre-treated with T0070907 prior to incubation (as indicated) with rosiglitazone and the lipid mix (n=1). **(E,F)** Relative mRNA expression levels in primary hepatocytes expressing Ad-**PGC-1α4** or Ad-Vector control pre-treated with T0070907 prior to incubation (as indicated) with glucagon (n=1). Data are expressed as means ± SD (n=1-2). *P ≤ 0.05, **P ≤ 0.01, ***P ≤ 0.001, and ****P ≤ 0.0001.

**SUPPLEMENTARY FIGURE 6.** ***The effect of modulating PGC-1α1 and PGC-1α4 expression on respiratory capacity of primary hepatocytes.***

**(A-L)** Basal, coupled, spare, non-mitochondrial, and maximal respiratory capacities, as well as proton leak, were assessed using oxygen consumption rate (OCR) measurements obtained from the Resipher System in primary hepatocytes with over-expressed and reduced PGC-1α1 and PGC-1α4 expression. Bars represent the mean ± SD of 6-14 biological replicate. *P ≤ 0.05, **P ≤ 0.01, ***P ≤ 0.001 and ****P ≤ 0.0001.

**SUPPLEMENTARY FIGURE 7.** ***Body composition and liver parameters of 12-week-old male and female PGC-1α4 liver transgenic mice after a 24-hour fasting period.***

**(A)** Liver acylglycerol levels, **(B,C)** liver weights, **(D)** serum triglyceride levels, **(E)** representative images of H&E stained of liver sections, **(F)** liver protein expression measured by Western blot **(G)** quantified by densitometry, and **H)** relative hepatic mRNA expression from 12-week-old $WT^{Cre+}$, $WT^{Tg+}$, and PGC-1α4$^{HepTg+}$ female mice following 24 hrs of fasting **(I)** Percentage of body weight loss in mice from A-F, calculated from values acquired before and after the 24-hr fast in each mouse. Bars are mean ± SEM (n=9-12 mice per group). *$P \leq 0.05$, **$P \leq 0.01$, ***$P \leq 0.001$, ****$P \leq 0.0001$.

**SUPPLEMENTARY FIGURE 8.** ***The effect of prolonged expression of* PGC-1α4 *on body weight and adipose tissue composition in male and female mic during fasting.***

Body weight (A,C) before and (B,D) after fasting, subqutenous (E,G) and visceral (F,H) adipose tissue weights from 12-week-old $WT^{Cre+}$, $WT^{Tg+}$, and PGC-1α$4^{HepTg+}$ female mice following 24 hrs of fasting. Bars are mean ± SEM (Males n=13, Females, n=9-12 mice per group). *$P \leq 0.05$, **$P \leq 0.01$, ***$P \leq 0.001$, ****$P \leq 0.0001$.

**SUPPLEMENTARY FIGURE 9.** ***The effect of reducing PGC-1α4 expression in the liver of male C57BL/6J mice on body composition, liver PPARγ protein and PPARγ target gene expression.***

**(A,B)** Fat and **(C,D)** lean mass (in grams and in %) measured by magnetic resonance imaging (MRI), **(E)** Relative protein expression levels of PPARγ measured by western blotting. **(F)** Relative mRNA expression levels of other PPARγ target genes (*Cd36, Adrp*) measured by qPCR in the livers of 12-week-old male AAV-shControl and AAV-shPGC-1α4 mice after 24 hours of fasting. Bars represent mean ± SEM for each cohort (n=9-11). *P ≤ 0.05, **P ≤ 0.01, ***P ≤ 0.001, ****P ≤ 0.0001.

**SUPPLEMENTARY FIGURE 10.** ***The effect of reducing PGC-1α4 expression in the livers of male C57BL/6J mice on glycolysis, glycogen synthesis and gluconeogenesis associated gene expression.***

**(A)** Glycolysis (*Gck, Pfkl, Eno1, Pklr*), **(B)** Glycogen synthesis (*Gys2*) and breakdown (*Pygl*) and **(C)** Gluconeogenesis (*Pepck, G6Pase*) associated gene expressions measured by qPCR in the livers of 12-week-old male AAV-shControl and AAV-shPGC-1α4 mice after 24 hours of fasting. Bars represent mean ± SEM for each cohort (n=9-11). (*P ≤ 0.05, **P ≤ 0.01, ***P ≤ 0.001, ****P ≤ 0.0001).

**SUPPLEMENTARY FIGURE 11.** ***Effect of PGC-1α4 expression modulation on glycolysis, glycogen synthesis & breakdown and gluconeogenesis-related gene expression in primary hepatocytes in-vitro.***

Relative mRNA expression of the genes involved in glycolysis (*Gck, Pfkl, Eno1, Pklr*) in primary hepatocytes **(A-D)** overexpressing control (Ad-Vector) or PGC-1α4 (Ad-PGC-1α4), or **(E-H)** silencing it by shRNA targeting control (shControl) and PGC-1α4 (shPGC-1α4). Glycogen synthesis (Gys2) and breakdown (*Pygl*) associated gene expressions) in primary hepatocytes **(I,J)** overexpressing control (Ad-Vector) or PGC-1α4 (Ad-PGC-1α4), or **(K,L)** silencing it by shRNA targeting control (shControl) and PGC-1α4 (shPGC-1α4). Glyconeogenesis (Pepck, G6Pase) associated gene expressions in primary hepatocytes **(M,N)** overexpressing control (Ad-Vector) or PGC-1α4 (Ad-PGC-1α4), or **(O,P)** silencing it by shRNA targeting control (shControl) and PGC-1α4 (shPGC-1α4) following 24-hour treatment with 50 nM glucagon or vehicle. Bars represent mean ± SD of technical replicates from a single experiment (n=4-8 independent biological replicates). *P ≤ 0.05, **P ≤ 0.01, ***P ≤ 0.001, ****P ≤ 0.0001.

**SUPPLEMENTARY FIGURE 12.** ***Histopathology assessment identifies steatosis, inflammation, hepatocellular ballooning and fibrosis stage in the livers of PGC-1α4 transgenic mice.***

**(A, B)** Steatosis level (as percentage) in the livers of 8-weeks old $WT^{Cre+}$, $WT^{Tg+}$, and PGC-$1α4^{HepTg+}$ male and female mice, respectively. Bars are mean ± SEM (Males, n=4-7; Females: n=3-7). *$P \leq 0.05$, **$P \leq 0.01$, ***$P \leq 0.001$, ****$P \leq 0.0001$. Pathology scores of **(C, E)** portal inflammation (0-1), **(D, F)** lobular inflammation (0-2), **(G, I)** hepatocellular ballooning and **(H, J)** fibrosis stages (0) in the livers of 8-weeks old (Males, n=4-7; Females: n=3-7).

**SUPPLEMENTARY FIGURE 13.** ***Body and liver parameters of PGC-1α4 liver transgenic female mice fed a HFHSHC diet.***

**(A)** Body weight, **(B) Fat and lean mass measured by MRI,** (C) Liver tissue weight, **(D)** serum triglycerides, **(E,F,J)** relative hepatic mRNA levels, **(G)** representative H&E stained liver sections, **(H)** liver acylglycerols, **(I)** serum ALT levels in female mice fed a high-fat, high-sucrose, high-cholesterol (Western) diet for 12 weeks. Data are expressed as mean ± SEM (n=10-11) ,*P ≤ 0.05, **P ≤ 0.01, ***P ≤ 0.001, ****P ≤ 0.0001.

**SUPPLEMENTARY FIGURE 14.** ***Effects of prolonged PGC-1α4 expression on liver injury in mice fed a HFHSHC Diet.***

**(A,F)** Relative mRNA expression levels in the livers of $WT^{Cre+}$, $WT^{Tg+}$, and PGC-1α$4^{HepTg+}$ male and female mice fed with HFHSHC diet for 12 weeks starting at 5 weeks of age. Data are shown as bar graphs of mean ± SEM (Males: n=11, females: n=10-11 per group). *P ≤ 0.05, **P ≤ 0.01, ***P ≤ 0.001, ****P ≤ 0.0001.

**SUPPLEMENTARY FIGURE 15.** Principal Component Analysis **(A)** score and **(B)** loading plot of untargeted lipidomic data collected from liver samples of PGC-1α$4^{HepTg+}$ and $WT^{Cre+}$ mice (n=43, male and female). Percentage of explained variance by PC1 and PC2 were 24.3% and 8.9% respectively.

**Table 1.** Primers applied for target gene amplification in the study

| Target gene (qPCR) | Forward primer (5'-3') | Reverse primer (3'-5') |
|---|---|---|
| **Mouse primers** | | |
| ***Acc1*** | CCAGCCTGAGCTACTCATAAA | CAAGAGGAGCTGGCAATACA |
| ***Aco2*** | GTTGGACCTCACCCAAAGAT | GGTCCGTGGTATTCCACAATAG |
| ***Acox1*** | CCCAAGACCCAAGAGTTCATT | GCCAGGACTATCGCATGATT |
| ***Acsl4*** | ACTAGGACCGAAGGACACATA | CCAATCCTACAGCCATAGGTAAA |
| ***Acta2*** | CCATCATGCGTCTGGACTT | GGCAGTAGTCACGAAGGAATAG |
| ***Acyl*** | CGGGAGGAAGCTGATGAATATG | GTCAAGGTAGTGCCCAATGAA |
| ***Adrp*** | ATTGCGGTTGCCAATACCTATGCC | TCGGACGTTGGCTGGTTCAGAATA |
| ***Arg1*** | GATTATCGGAGCGCCTTTCT | TGGTCTCTCACGTCATACTCT |
| ***Atgl*** | CATGATGGTGCCCTATACTCTG | CTACCCGTCTGCTCTTTCATC |
| ***Atp5e*** | CTACTCTGAAGCGACCCAGC | GGGAAAACCGGATGTAGCTGA |
| ***Atpsynf1*** | TCTCCATGCCTCTAACACTCG | CCAGGGTCAACAGACGTGTCAG |
| ***B220*** | AATGGAGACCAGGAAGTCTGTGCT | TGTAGGCTGAGGCTCTGTTTGTGT |
| ***Catalase*** | GCGGATTCCTGAGAGAGTGGTAC | GCCTGACTCTCCAGCGACTGTGGAG |
| ***Cd11b*** | TTACCTGGGTTATGCTTCTGCAG | AAGCTTTGGACACGGTTCCTC |
| ***Cd11c*** | GTGCTGAGTTCGGACACAGT | AGAGGCCACCTATTTGGTTAGT |
| ***Cd36*** | CTGGGACCATTGGTGATGAAA | CACCACTCCAATCCCAAGTAAG |
| ***Cd4*** | ACCTCAAGCTCCAGCTGAAGGAAA | GGTTGCCAGAACCAGCAAACTGAA |
| ***Col1a1*** | GAAACCCGAGGTATGCTTGA | GTTGGGACAGTCCAGTTCTT |
| ***Cox4i*** | AGTTTAACGAGAGCTTCGCCGAGA | AGCGCAGTGAAGCCAATGAAGAAC |
| ***Cox5b*** | ATGCTACCTCCAAAGGCAGCTTC | TGCAGCCCACTATTCTCTTGTTGC |
| ***CoxI*** | ACTTGCAACCCTACACGGAGGTAA | TCGTGAAGCACGATGTCAAGGGAT |
| ***CoxIII*** | GCAGGATTCTTCTGAGCGTTCT | GTCAGCAGCCTCCTAGATCATGT |
| ***Cpt1a*** | GAACCCCAACATCCCCAAAC | TCCTGGCATTCTCCTGGAAT |
| ***Cs*** | CAAGCAGCAACATGGGAAGA | GTCAGGATCAAGAACCGAAGTCT |
| ***Cyp4a10*** | CCTACATCTCAAGAAGCTCCAC | GGTAGGTCACACCAACAACTTA |
| ***Cyp4a14*** | TCTCATCTTTCTGCCCTCATTTC | CAGTGGCTGGTCAGAGTTAAAG |
| ***Cyp4a31*** | GGGTATGGTTTGCTCCTGTTA | TCAGAATGTCATAGTGGAAGGC |
| ***Cyp4f14*** | AGGAACTATCGTCACCTCCA | GTACCCACCAAACGAGTCAA |
| ***Cyp4f18*** | GGCAAGACACCCAGAATACC | CAGGTCGTCCCATTCAATCTC |
| ***Cyto-b*** | ACAAACCTCCTATCAGCCATCCCA | AGTGGAAAGCGAAGAATCGGGTCA |
| ***Cyto-c*** | GCAAGCATAAGACTGGACCAAA | TGTTGGCATCTGTGTAAGAGAATC |
| ***Eno1*** | CTACAGGTCTGGCAAGTATGAC | GTAGTTCTGGACGAAGGACTTG |
| ***F4/80*** | CTTTGGCTATGGGCTTCCAGT | GCAAGGAGGACAGAGTTTATCGTG |
| ***Fasn*** | TCCTGGAACGAGAACACGATCT | GAGACGTGTCACTCCTGGACTTG |
| ***Fsp27/Cidec*** | CGTGTTAGCACCGCAGAT | AAGATGTCCTGGACCTTGTTC |
| ***G6Pase*** | CAGGCATTGCTGTGGCTGAAACTT | TAGCAGGTAGAATCCAAGCGCGAA |
| ***Gck*** | CAGGCTGACACCCAACT | CACCGCAGAGACCAGTG |
| ***Glut2*** | AATGTACTGGAAGCAGAGGGCGAT | AGGTCCAATCCCTTGGTTCATGGT |
| ***Gpx1*** | CAGGAGAATGGCAAGAATGAAGAG | GGCATTCCGCAGGAAGGTAAAGAGC GG |
| ***Gys2*** | CTGATGCAGACCAACAGATA | CCGTTACAGTAAGGTGACTC |

| | | |
|---|---|---|
| ***Hmgb1*** | ATGGGCAAAGGAGATCCTA | ATTCATCATCATCATCTTCT |
| ***Hprt*** | GGCCAGACTTTGTTGGATTTG | TGCGCTCATCTTAGGCTTTGT |
| ***Hsl*** | CATCAACCACTGTGAGGGTAAG | AAGGGAGGTGAGATGGTAACT |
| ***Il-10*** | GCTCTTACTGACTGGCATGAG | CGCAGCTCTAGGAGCATGTG |
| ***Il-1β*** | AAGGGCTGCTTCCAAACCTTTGAC | ATACTGCCTGCCTGAAGCTCTTGT |
| ***Lcad*** | CTTGCTTGGCATCAACATCGCAGA | ATTGGAGTACGCTTGCTCTTCCCA |
| ***Ly6c1*** | GCAGTGCTACGAGTGCTATGG | ACTGACGGGTCTTTAGTTTCCTT |
| ***Mcad*** | AACACTTACTATGCCTCGATTGCA | CCATAGCCTCCGAAAATCTGAA |
| ***Mcp1*** | TGCTGTCTCAGCCAGATGCAGTTA | TACAGCTTCTTTGGGACACCTGCT |
| ***Mmp9*** | TCTGTATGGTCGTGGCTCTAA | GGAGGTATAGTGGGACACATAGT |
| ***Mrc1*** | CTCTGTTCAGCTATTGGACGC | CGGAATTTCTGGGATTCAGCTTC |
| ***Ndufs1*** | AGGATATGTTCGCACAACTGG | TCATGGTAACAGAATCGAGGGA |
| ***Ndufs5*** | GACATACAGAAAAAGCTGGGCA | TCGCCTCATCGTTTTGTACCG |
| ***Pdhae1*** | GCTGGCATAAACCCTACGGAC | CCTTTCCCTTTAGCACAACCTC |
| ***Pepck*** | CAGGATCGAAAGCAAGACAGT | AAGTCCTCTTCCGACATCCAG |
| ***Pgc-1a1*** | GGACATGTGCAGCCAAGACTCT | CACTTCAATCCACCCAGAAAGCT |
| ***Pgc-1a4*** | TCACACCAAACCCACAGAAA | CTGGAAGATATGGCACAT |
| ***Pparg*** | CAAGAATACCAAAGTGCGATCAA | GAGCTGGGTCTTTTCAGAATAATAAG |
| ***Prx5*** | GGGAAGGCGACAGACTTATTATT | CCTTCACTATGCCGTTGTCTATC |
| ***Pygl*** | CTGTGGCAGAAGTGGTGAA | GATAGGTCTGTGGCTGGAATG |
| ***Scad*** | ACCAAAGCTTGGATCACCAACTCC | AACCAGGAAGGCACTGATACCCTT |
| ***Scd1*** | ACAGCCTGTTCGTTAGCACCTTCT | CCCGGGATTGAATGTTCTTGTCGT |
| ***Sdh*** | GTTGGCGCAGTTTCGAGGCT | GCCGCAGGTCTGTTTTTGGA |
| **Srebp1c** | CATCGACTACATCCGCTTCTT | CACCAGGTCCTTCAGTGATTT |
| ***Sod2*** | GGCCAAGGGAGATGTTACAAC | GCAACTCTCCTTTGGGTTCTC |
| ***Tgfβ1*** | GCAACAATTCCTGGCGTTAC | GTATTCCGTCTCCTTGGTTCAG |
| ***Tnfα*** | AGCCGATGGGTTGTACCTTGT | TGAGATAGCAAATCGGCTGAC |
| ***Uqcrc1*** | TGCCTTAGAGAAGGAGGTAGAG | GACAGTGCCTTGATGAGGTAAG |
| **Vlcad** | GGCCAAGCTGGTGAAACACAAGAA | ACAGAACCACCACCATGGCATAGA |
| **Ym1** | ATCTATGCCTTTGCTGGAATGC | TGAATGAATATCTGACGGTTCTGAG |
| **Target gene (qPCR)** | **Forward primer (5'-3')** | **Reverse primer (3'-5')** |
| **Human primers** | | |
| ***PGC-1α1*** | CCTTATTTTCTCAAAGACCC | GGATCTTGAAGAGGATCTAC |
| ***PGC-1α4*** | TCACACCAAACCCACAGA | CTGGAAGATATGGCACAT |
| **Proximal Promoter (Exon-1a)** | GGACATGTGCAACCAGGACT | CACTTGAGTCCACCCAGAAAGCT |
| **Alternative promoter (Exon-1b)** | GCACTGCAGTAAAATGAATGACAC | GTTCAGGAAGATCTGGGCAAA |
| **Alternative promoter (Exon-1b')** | GTGAGTATCAGGAGGCATTATG | TTCAGGAAGATCTGGGCAAAGA |
| ***HPRT*** | CCTGGCGTCGTGATTAGTGAT | AGACGTTCAGTCCTGTCCATAA |
| **Target gene (non-qPCR)** | **Forward primer (5'-3')** | **Reverse primer (3'-5')** |
| ***PGC-1α1*** | GACATGTGCAACCAGGACTC | CTCAAATGGGGAACCCTTGG |
| ***NT-PGC-1α-a*** | GACATGTGCAACCAGGACTC | CTGGAAGATATGGCACAT |
| ***PGC-1α4*** | GGTTATCATCTATGGATTC | CTGGAAGATATGGCACAT |

| | | |
|---|---|---|
| ***PGC-1α-b*** | GGTTATCATCTATGGATTC | GTCACGTCTCCATCTGTCAG |
| ***PGC-1α-c*** | CTAGGAGGCTTTATGCTATTG | CTGGAAGATATGGCACAT |
| ***L-PGC-1α*** | GGTAGCAAGATTTATGTTTC | GAGGGCAATCCGTCTTCATCC |
| ***18S*** | CTCAACACGGGAAACCTCAC | CGCTCCACCAACTAAGAACG |
